# MAERM: Predicting Enzyme-Reaction Matching Relationships with a Mixed-Attention Model

**DOI:** 10.64898/2026.07.06.736902

**Authors:** Tiantao Liu, Silong Zhai, Shaolong Lin, Xinke Zhan, Junwen Deng, Huanxiang Liu, Shirley W. I. Siu

**Affiliations:** Faculty of Applied Sciences, Macao Polytechnic University, 999078, Macau SAR, China; College of Pharmaceutical Sciences, Zhejiang University, Hangzhou 310058, Zhejiang, China

**Keywords:** enzyme-reaction matching, mixed-attention model, enzyme promiscuity, reaction center, chemoenzymatic catalysis

## Abstract

Harnessing enzyme specificity requires a thorough understanding of enzyme promiscuity, which determines enzymes’ catalytic scope; however, measuring this scope still relies heavily on labor-intensive analytical approaches. While data-driven approaches have emerged to predict the catalytic scope of enzymes, these methods continue to face challenges such as restricted datasets and insufficient integration of enzyme structural information and reaction transformations. Here, we introduce MAERM, an innovative mixed-attention model designed to predict enzyme-reaction matching relationships. Built on our MAERM-DB, a dataset with broad coverage of validated and chemoenzymatic catalysis data, MAERM utilizes a local-global attention module to integrate multimodal enzyme information with fine-grained reaction representations, thereby predicting enzyme-reaction matching probabilities. Results show that MAERM consistently outperforms all baselines, with an average F1-score of 0.984. Notably, on challenging test samples with less than 40% sequence identity to the training set, MAERM outperforms the second-ranked model by 5.9% in F1-score. In addition, MAERM achieves the highest top-10 success rate of 51.7% on Enzyme-405 and the highest balanced accuracy of 0.697 on BioCat-547, further supporting its generalizability in enzyme screening and chemoenzymatic catalysis. Finally, MAERM can serve as an efficient scoring module. When integrated with ProteinMPNN, MAERM has successfully guided novel enzyme design for two carbonyl reduction reactions, resulting in enhanced catalytic potential for the native substrate and demonstrating broad compatibility. Overall, MAERM has the potential to reduce the experimental cost of measuring enzymes’ catalytic scope, facilitate enzyme design, and ultimately accelerate the design-build-test-learn cycle in enzyme engineering.

## 1. Introduction

Enzymes play a pivotal role in drug discovery and agriculture, serving as mild, sustainable, and efficient catalysts for biotransformations [1]. Rapid advances in enzyme discovery and engineering, including directed evolution and cascade catalysis [2], have accelerated their adoption in large-scale biocatalytic processes. For instance, the one-pot enzymatic cascade enables the production of Islatravir (an HIV prophylactic) with multiple chiral centers from a single reaction vessel and basic building blocks (e.g., alkynyl glycerol) [3]. Enzyme promiscuity is a key determinant of biocatalytic applications, as it allows enzymes to catalyze reactions beyond their native functions and thereby expands their catalytic scope [4].

Enzyme promiscuity shapes biocatalytic utility in complex and sometimes opposing ways. Enabling catalytic promiscuity to overcome enzyme specificity for particular reactions is arguably the most significant strategy for expanding catalytic scope [5]. Yet, a high level of promiscuity is often associated with unwanted side effects, such as compromised biological function and poor catalytic properties [6]. For instance, enzymes with high promiscuity often exhibit low catalytic efficiency for reactions they catalyze best [7]. Measuring the extent of promiscuity, i.e., the catalytic scope of an enzyme, helps define its utility. In biocatalysis, expanding natural enzymes to perform non-native transformations (i.e., chemoenzymatic catalysis) is an important objective [8, 9]. Thus, precise measurement of enzymes’ catalytic scope becomes essential for defining their functional limits and guiding the discovery of new reaction types [10].

Researchers can retrieve catalytic information from biological databases such as BRENDA, KEGG, MetaCyc, and Rhea [11–14]. However, reactions recorded in these databases represent merely a small fraction of an enzyme’s actual catalytic scope. This gap is evident in MetaCyc, which assigns only 20,039 reactions to 14,473 enzymes [13], whereas current estimates suggest that each enzyme can catalyze an average of approximately ten reactions [15]. Moreover, these databases primarily focus on metabolic pathways and therefore capture relatively little chemoenzymatic catalysis, further limiting their applicability. As a result, measuring the catalytic scope of enzymes still relies on labor-intensive analytical approaches, such as gas chromatography-mass spectrometry (GC-MS) [16, 17]. These manual analyses are inefficient, costly, and difficult to scale to the large numbers of mutants generated by directed evolution [18]. Data-driven approaches are therefore urgently needed to address these limitations.

Data-driven approaches have been developed to predict enzyme substrate specificity (i.e., whether an enzyme can catalyze given substrates), but most efforts have focused on specific protein families [19–21]. ESP applies a graph neural network (GNN) to model substrate specificity for more general enzymes [22], but still shows limited coverage of non-native substrates. To address this gap, EZSpecificity introduces the ESIBank dataset [23], which integrates a broad spectrum of native and non-native substrates with sequence-, structure-, and interaction-level information. The model also adopts an SE(3)-equivariant GNN architecture to capture local atomic environments at catalytic sites, achieving improved performance compared with ESP. Although these models enable rapid substrate-specificity prediction, they rely only on substrate information and do not account for other reaction components. Products play an indispensable role in enzymatic reactions, as their release may constitute the rate-limiting step of the catalytic process. For instance, in the deamination of cytosine to uracil catalyzed by yeast cytosine deaminase (yCD), cleavage of the uracil-metal bond and subsequent product release are rate-limiting steps [24]. Therefore, reliable prediction of catalytic scope requires consideration of both reactants and products, which are essential for representing the full biological function of enzymes [25]. Several recent models have extended prediction from substrate specificity to enzyme-reaction matching [26–28]. These models take a full reaction and a candidate enzyme as input to assess whether the enzyme can catalyze the reaction. While these models provide frameworks for integrating reaction and enzyme data, they remain limited by three primary constraints: (1) their datasets are confined to metabolic pathways, restricting their applicability to chemoenzymatic catalysis; (2) they rely predominantly on sequence data, without fully incorporating multimodal enzyme features; and (3) they encode entire reactions, lacking sensitivity to local reactive regions.

In this work, we introduce MAERM, a mixed-attention model designed to predict enzyme-reaction matching relationships. We first construct MAERM-DB, a dataset that integrates enzyme-reaction pairs derived from ESP [22], the Mechanism and Catalytic Site Atlas (M-CSA) [29], and RetroBioCat-DB (RDB) [30]. Compared with existing datasets [27, 31, 32], MAERM-DB is designed to provide broader coverage of validated and chemoenzymatic catalysis data. Previous datasets curate enzyme-reaction pairs by retrieving enzymes based on the reactions’ EC numbers. However, a specific EC number can correspond to a large and highly redundant enzyme set. For example, more than 230,000 enzyme sequences in Swiss-Prot are annotated under EC 1.1.1.2 [33]. These datasets often lack effective filtering of redundant enzymes, as enzymes sharing the same EC number may still differ in their substrate preferences [34]. In contrast, MAERM-DB does not retrieve enzyme-reaction pairs based on EC numbers. Instead, it integrates pairs supported by literature evidence from M-CSA and by experimental validation or phylogenetic confirmation from ESP. Incorporating RDB further expands its coverage of chemoenzymatic reactions and supports broader applicability.

Built on MAERM-DB, we propose a binary classification model to predict enzyme-reaction matching relationships. The architecture consists of three components: (1) the enzyme encoder (Figure 1a), generating enzyme embeddings by integrating the features of ESM Cambrian (ESM-C) [35] and the geometry-sequence graph [36] through a cross-attention module; (2) the reaction encoder (Figure 1b), generating reaction embeddings from the local reactive region (i.e., reaction centers) and the global reaction context (i.e., the whole reaction); and (3) the local-global attention module (Figure 1c), integrating enzyme embeddings with both the local reactive region and the global reaction context to predict enzyme-reaction matching probabilities.

**Figure 1.**
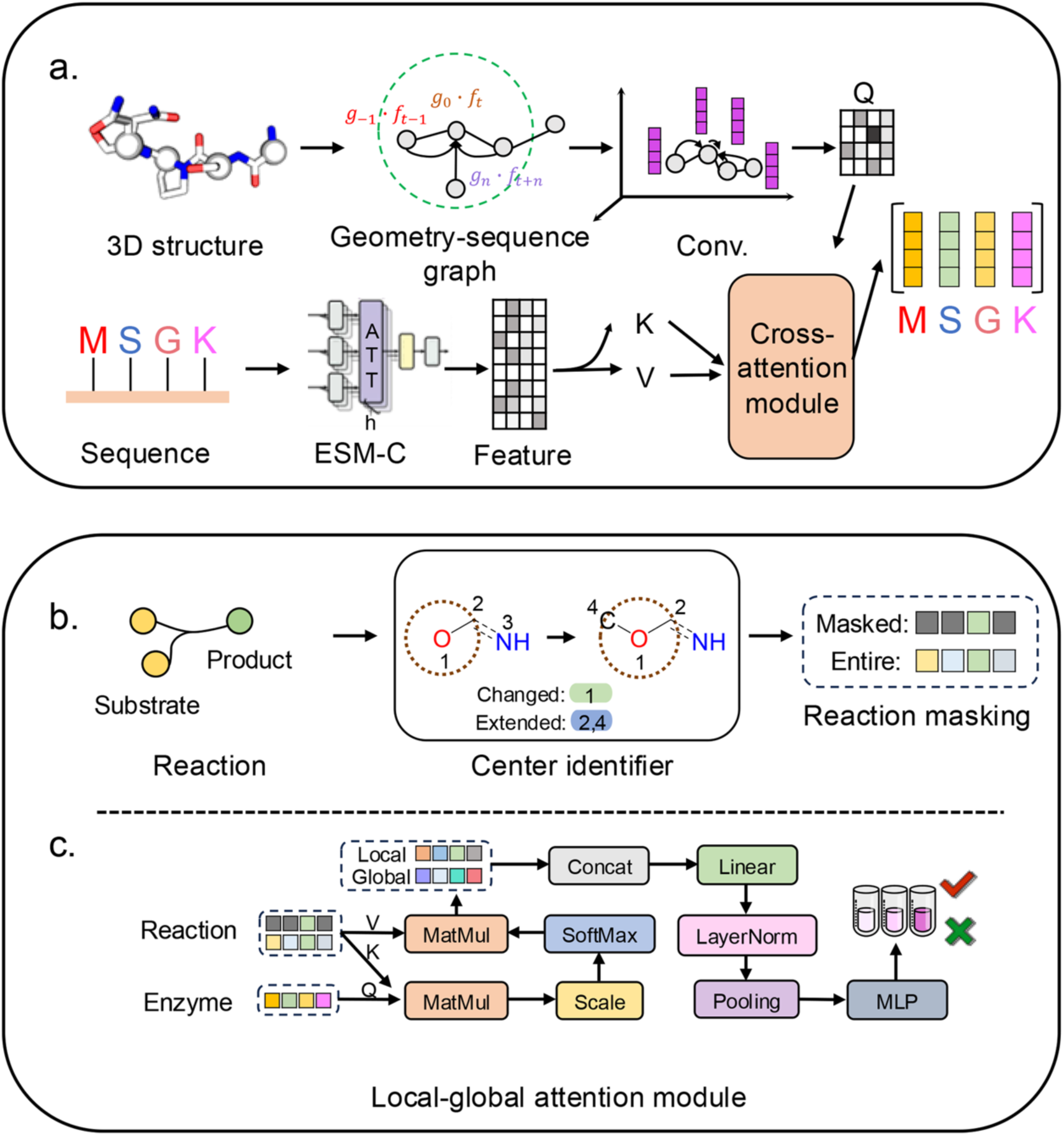
The architecture of our model. (a) The enzyme encoder generates enzyme embeddings by integrating the features of ESM-C and the geometry-sequence graph through a cross-attention module. (b) The reaction encoder generates reaction embeddings from the local reactive region and the global reaction context. (c) The local-global attention module integrates enzyme embeddings with both the local reactive region and the global reaction context to predict enzyme-reaction matching probabilities.

Compared with previous approaches [26–28], our work makes the following key contributions: (1) The introduction of the geometry-sequence graph and ESM-C enables MAERM to capture multimodal enzyme information; (2) The local-global attention module integrates enzyme embeddings with reactive regions and global reaction context, providing a careful adaptation of fine-grained reaction representations; (3) Baseline comparisons across models for drug-target interaction (DTI) prediction, enzyme substrate specificity prediction, and enzyme-reaction matching prediction provide a comprehensive evaluation; (4) Interpretability analyses using perturbation-based methods reveal key residues underlying real binding. Results show that MAERM consistently outperforms all baselines, with an average F1-score of 0.984. Notably, on challenging test samples with less than 40% sequence identity to the training set, MAERM outperforms the second-ranked model by 5.9% in F1-score. In addition, MAERM achieves the highest top-10 success rate of 51.7% on Enzyme-405 and the highest balanced accuracy of 0.697 on BioCat-547, underscoring its utility in enzyme screening and chemoenzymatic catalysis. Finally, MAERM can serve as an efficient scoring module when integrated with the protein design tool ProteinMPNN [37] to guide enzyme design for two carbonyl reduction reactions. Overall, MAERM provides a new perspective on enzyme-reaction matching prediction and may help reduce the experimental cost of measuring enzyme catalytic scope, advance enzyme design, and accelerate the design-build-test-learn cycle in enzyme engineering.

## 2. Methods

### Original data curation

A benchmark dataset, MAERM-DB, was curated from three independent sources (Figure 2a). The first source is the ESP dataset [22], which contains enzyme-substrate pairs. We extended the original pairs to enzyme-reaction pairs using experimentally or phylogenetically confirmed gene ontology (GO) annotations [38] provided in ESP. Each GO annotation links the UniProt IDs [39] of enzymes and the ChEBI IDs [40] of all compounds in the reaction. Thus, we obtained enzyme sequences and reaction SMILES through these IDs. In this study, we name the extended ESP dataset as ESP-ER to distinguish it from the original ESP dataset. The second source is the Mechanism and Catalytic Site Atlas (M-CSA) dataset [29], which catalogs reaction mechanisms and catalytic sites. We extracted enzyme sequences and reaction SMILES based on the associated UniProt IDs and ChEBI IDs. The third source, RetroBioCat-DB (RDB) [30], focuses on chemoenzymatic reactions involving organic substrates. The enzyme-reaction pairs were obtained via RDB’s search portal for exploring substrate specificity. For each enzyme, we retrieved the images of corresponding substrates and products. Then, we used img2mol [41] to convert these images to SMILES, thereby assembling the enzyme-reaction pairs. Following the preprocessing protocol of the ECREACT dataset [42], common coenzymes and byproducts (Table S1) were excluded from the reactions in MAERM-DB. For computational efficiency, we further limited SMILES strings to a maximum length of 500 and protein sequences to 1,000 residues.

**Figure 2.**
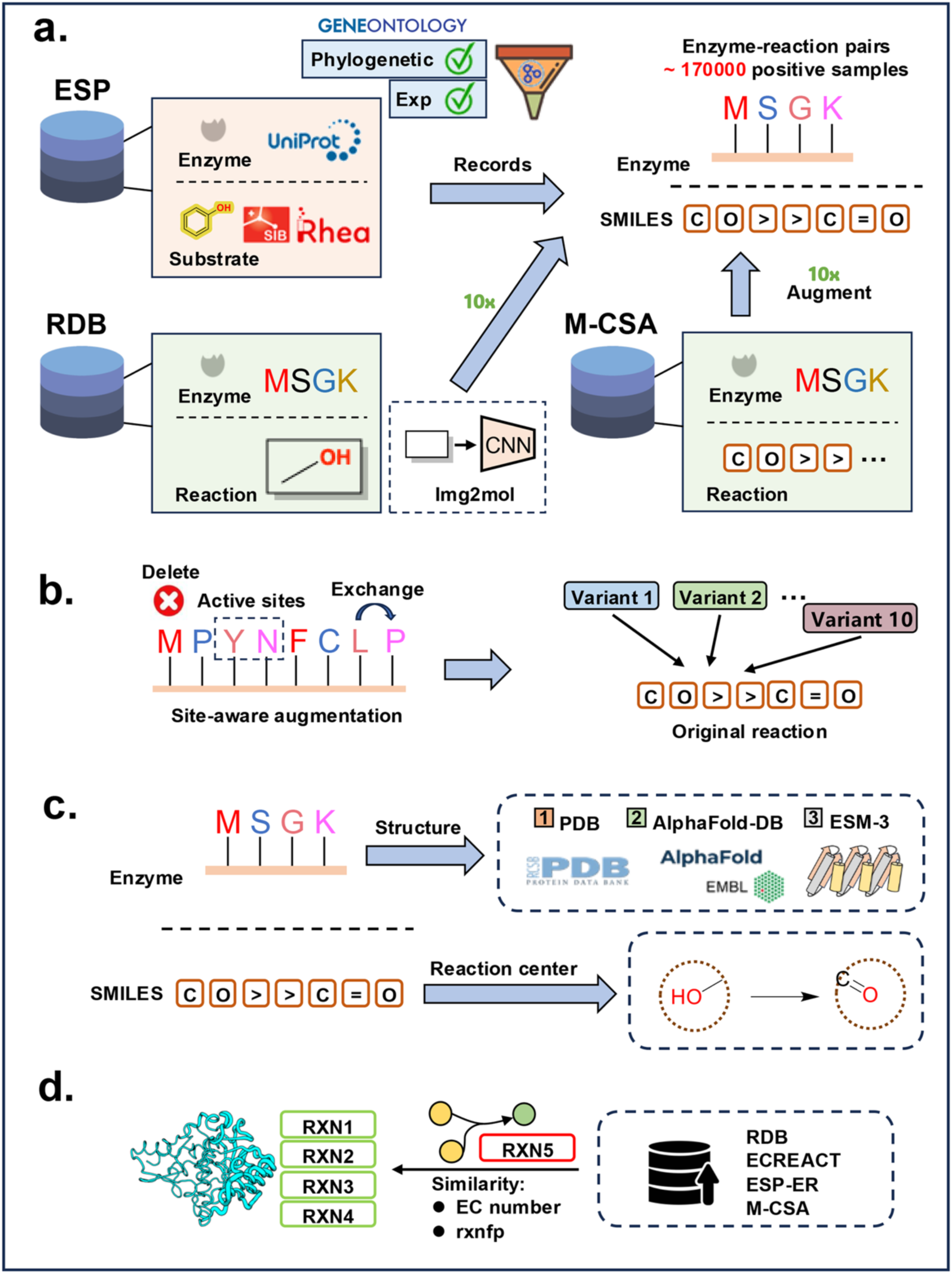
Overview of the curation of MAERM-DB. (a) Original data curation for ESP-ER, RDB, and M-CSA. (b) Site-aware augmentation applied to the training sets of M-CSA and RDB. (c) Information extraction for enzymes and reactions. (d) Negative sample generation based on similarity.

### Data augmentation and information extraction

We applied augmentation to generate plausible enzyme variants for the training sets of M-CSA and RDB to balance data. Inspired by previous work [32, 43], we utilized site-aware augmentation (Figure 2b). We first augmented the enzyme sequences by 10-fold through four basic operations: random substitution, insertion, deletion, and swap (Table S2). During this process, annotated sites from UniProt (i.e., binding sites, active sites, and sites) or M-CSA were preserved whenever available.

As shown in Figure 2c, the structures of all enzymes were obtained in the following order of priority: if a PDB ID [44] was available, the reported crystal structure was used; if only an AlphaFold-DB (AFDB) ID [45] was available, the AlphaFold-predicted structure was used; and if only the sequence was available, the structure was predicted using ESM3-small (version 2024-08) [46]. For all reactions, atom mapping was computed using RXNMapper [47], and RDChiral_cpp [48] was subsequently used to extract reaction centers within fixed radii of 0, 1, and 2. These reaction centers were anchored by the atoms directly involved in the reaction, and the corresponding local regions were defined by expanding from these atoms to the specified radius.

After removing enzyme-reaction pairs with failed information extraction, we employed CD-HIT [49], a widely used greedy incremental algorithm for clustering protein sequences. We applied a cutoff of 0.8 to ensure that the training, validation, and test sets shared less than 80% sequence identity. As shown in Table S3, the positive samples consisted of 140,128 training samples (with data augmentation), 14,050 validation samples, and 13,598 test samples.

### Negative pair generation

For negative enzyme-reaction pairs, we noted that studies of enzyme catalytic scope typically test chemically similar substrates [16, 17]. For instance, substrate scope experiments for 4-phenol oxidases often aim to identify a variety of phenolic substrates with substituents at p- and o-position to the hydroxy group [16]. Thus, as shown in Figure 2d, we generated negative samples by replacing the reaction in each positive pair with a similar reaction (i.e., a negative reaction). Negative reactions were selected from a larger reaction dataset that integrates enzymatic reactions from ESP-ER, M-CSA, RDB, and ECREACT. Specifically, for each positive reaction, we selected the most similar but not identical reaction with the same EC number as the negative, based on the cosine similarity of rxnfp embeddings [50]. For positive reactions lacking an EC number, the negative sample was derived by selecting the reaction with the highest similarity from the entire dataset. All negative samples were then processed using the same data augmentation, information extraction, and splitting procedures as the positive samples, resulting in 122,759 training samples, 13,722 validation samples, and 13,263 test samples. Using CD-HIT, we grouped the test set of MAERM-DB into three sequence identity levels relative to the training data (0–40%, 40–60%, and 60–80%), yielding 3,252, 11,117, and 12,492 samples, respectively (Table S4).

### Dataset analysis

To investigate the distributional landscape of MAERM-DB, we employed TMAP [51]. Reaction representations were computed using rxnfp, while enzyme representations were obtained using ESM-C (600M, version 2024-12). Enzyme-reaction pair features were constructed by concatenating the ESM-C and rxnfp representations, and all features were subsequently projected into a low-dimensional space using TMAP. Moreover, AutoDock Vina (v1.2.5) [52] was employed to further characterize the differences between positive and negative enzyme-reaction pairs. We performed docking for all positive and negative pairs in the RDB dataset and calculated the corresponding average Vina scores. To leverage available active-site information, a docking box with a radius of 20 Å centered on the enzyme active site was defined. All substrates from a single reaction were docked independently within the same box using an exhaustiveness parameter of 8. The final average Vina score for an enzyme-reaction pair was defined as the average of the best Vina scores across all associated substrates.

### Enzyme Encoder

The first component of MAERM is the enzyme encoder (Figure 1a). The encoder integrates features from ESM-C (600M, version 2024-12) and the geometry-sequence graph [36] through a cross-attention module. The geometry-sequence graph represents an enzyme as a sequence of features (*f*_1_, *f*_2_, …, *f_n_*), where *f_t_ε*ℝ^c×1^ denotes the feature of the *t*-th amino acid. Each amino acid is also associated with a 3D coordinate *p_t_ε*ℝ^3×1^, extracted from the enzyme’s corresponding PDB file. The feature *f_t_* is subsequently updated via a (3+1)D convolution kernel:

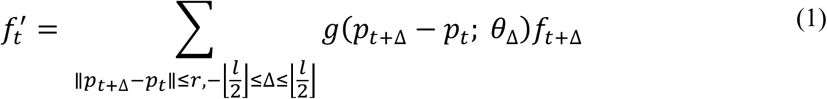

where *r* and *l* are the geometric radius and the sequential kernel size, *g*(·) and *θ*_Δ_ are the kernel function and weights, respectively. This equation produces *E*_graph_ for each enzyme.

The (*l* + 1)*^t^*: layer of the cross-attention module [53] takes the outputs *E*_graph_ = [*g*_1_, *g*_2_, …, *g_n_*] and *E*_ESM-C_ = [*c*_1_, *c*_2_, …, *c_n_*]. The attention components are computed as follows:

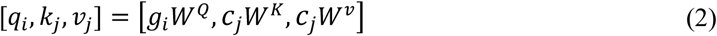

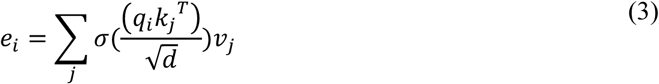

where σ is the SoftMax operation, *d* is defined as *d*_*esm*/h, where the ESM-C dimension *d*_*esm* is 1152 and the number of attention heads h is set to 8 during training. The details of enzyme encoder are provided in **Supporting Information Part 1**.

### Reaction encoder

The second component of MAERM is the reaction encoder (Figure 1b). A reaction SMILES string can be denoted as *R* = [*r*_1_, *r*_2_, *r*_’_, …, *r_n_*]. As described above, RDChiral_cpp [48] was used to automatically extract reaction centers with radii of 0, 1, and 2. The reaction center can be represented as a subset *R_rc_* ⊂ *R*, which contains only the tokens within the center, while all tokens outside this region are masked. Inspired by a previous study [54], we utilized an annealing strategy to represent the balanced reaction center *R*⎯. Specifically, the annealing strategy initially favors *R* with high probability and gradually shifts toward *R_rc_* in the later stages:

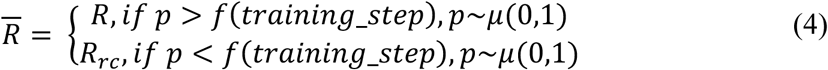

Here, *μ* denotes a value randomly sampled from the interval [0,1]. *f*(*training*_*step*) is defined as a monotonically increasing concave function over the training process: 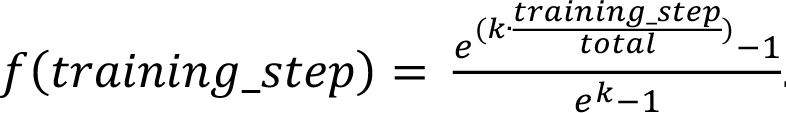. In our setup, *total* (i.e., total training steps) and *k* are set as 8,215,000 and 2, respectively.

### Local-global attention module

The local-global attention module (Figure 1c) takes the outputs from previous encoders: the enzyme encoder output *E*, and the reaction encoder outputs *R* and *R*⎯. We implement a mixed local-global attention mechanism to integrate the information from enzyme and reaction. For the *i^th^* token of enzyme sequence, and the *j^th^* token of reaction SMILES, the corresponding features in E, *R*, and *R̅* are denoted as *e_i_*, *r_j_*, and *r̅_j_*, respectively. The (*l* + 1)*^th^* layer of the local-global attention module *g^l^*^+1^, is defined as follows:

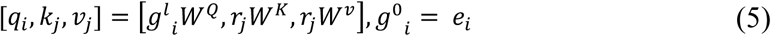

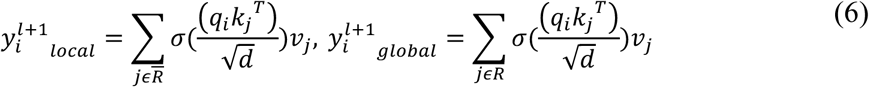

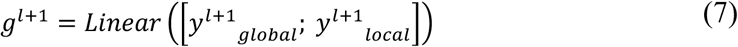

where *σ* denotes the SoftMax operation, and *d* is defined as *d*_*model*/h. The model dimension *d*_*model* was set to 256 and the number of attention heads h was set to 8 during training. The local-global attention module consists of 8 layers, and its output is aggregated across all nodes using mean pooling.

A multi-layer perceptron (MLP) classifier was utilized to map the 256-dimensional representations to the binary classes. Cross-entropy loss was used between the predicted probabilities and the labels. Models were trained using the stochastic gradient descent (SGD) optimizer, with three independent runs under different seeds. Details of the local-global attention module are provided in **Supporting Information Part 2**.

### Ablation studies

Extensive ablation experiments were carried out to examine the contribution of different model components and training strategies (as shown in **Table 1**). In the enzyme encoder, we evaluated models without the geometry-sequence graph (MAERM-w/o Graph), without ESM-C features (MAERM-w/o Seq), and with ESM-C features replaced by residue-level one-hot embeddings (MAERM-Residue). For the reaction encoder, we trained models without the annealing strategy (MAERM-SMILES and MAERM-Center). In these two settings, the reaction representation *R⎯* in Equation 4 is fixed to the full reaction R or reaction center *R_rc_* throughout training, respectively. Moreover, we examined the effect of reaction center radius. The default radius of 1 in MAERM was replaced with radii of 0 and 2, resulting in MAERM-R0 and MAERM-R2. Except for the specified modifications, all other components of the models were identical to those of MAERM.

**Table 1.**
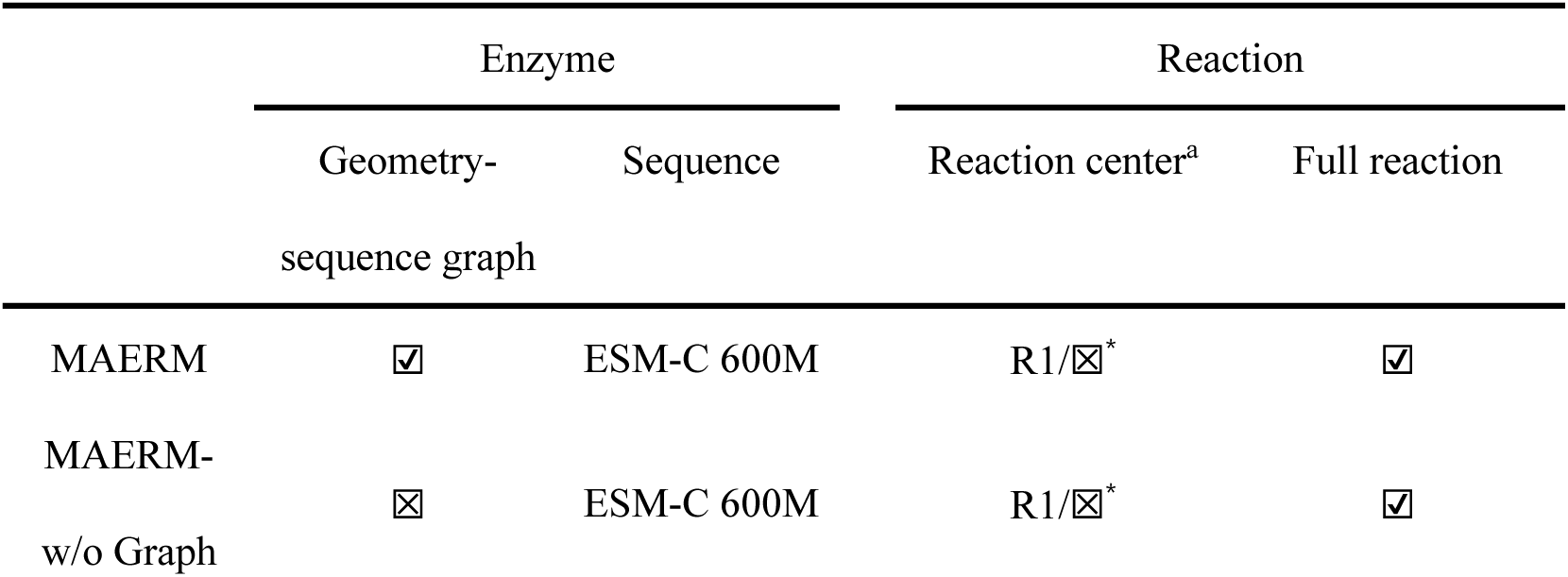
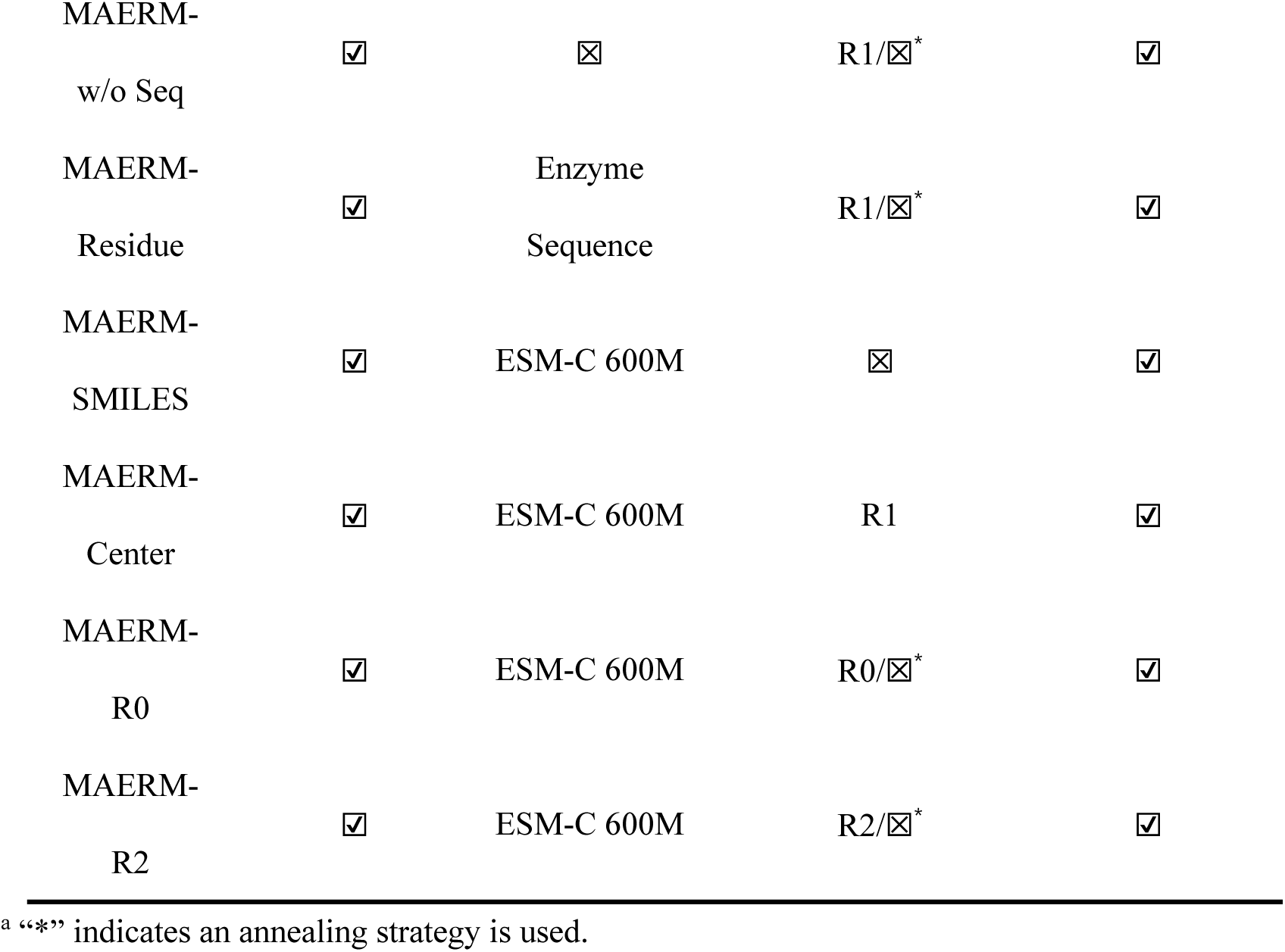
Experimental settings for MAERM ablation studies.

|  | Enzyme |  | Reaction |  |
| --- | --- | --- | --- | --- |
|  | Geometry-sequence graph | Sequence | Reaction center<sup>a</sup> | Full reaction |
| MAERM | ☑ | ESM-C 600M | R1/☒<sup>*</sup> | ☑ |
| MAERM-w/o Graph | ☒ | ESM-C 600M | R1/☒<sup>*</sup> | ☑ |
| MAERM-w/o Seq | ☑ | ☒ | R1/☒* | ☑ |
| MAERM-Residue | ☑ | Enzyme<br>Sequence | R1/☒* | ☑ |
| MAERM-<br>SMILES | ☑ | ESM-C 600M | ☒ | ☑ |
| MAERM-Center | ☑ | ESM-C 600M | R1 | ☑ |
| MAERM-R0 | ☑ | ESM-C 600M | R0/☒* | ☑ |
| MAERM-R2 | ☑ | ESM-C 600M | R2/☒* | ☑ |

<sup>a</sup> “\*” indicates an annealing strategy is used.

### Baseline model

Baseline models were selected from diverse sources, including SPEPP [26], CACLENS [27], ESP [22], MCANet [55], ML-DTI [56], MAARDTI [57], and Rep-ConvDTI [58]. Among these, SPEPP and CACLENS are available for predicting enzyme-reaction matching relationships. Due to the training script availability, we adopted CACLENS-TU, which is built upon ProteinT5 and UniMol [59, 60]. ESP utilizes a gradient boosting model to predict enzyme substrate specificity. In this study, the original substrate SMILES were replaced with reaction SMILES to adapt to our task. MCANet, ML-DTI, MAARDTI, and Rep-ConvDTI are originally developed for drug-target interaction (DTI) prediction using protein sequences and drug SMILES. Similarly, drug SMILES were replaced with reaction SMILES. In the reproduction setting, the main hyperparameters were kept the same as in the original models. All baseline models were retrained three times on the MAERM-DB dataset using a single NVIDIA A800 GPU. More details are provided in **Supporting Information Part 3**.

### Evaluation metrics and interpretability

The performance of MAERM was evaluated using standard metrics, including accuracy (Acc), recall, precision, F1-score, area under the receiver operating characteristic curve (AUROC), and area under the precision-recall curve (AUPR). All standard metrics were computed using Scikit-learn [37]. The definitions are provided in **Supporting Information Part 4**.

The interpretability of MAERM was examined from both the enzyme and reaction sides. For enzyme-side interpretability, we employed a perturbation-based analysis inspired by previous work [61]. We defined the contribution of each residue to the overall prediction as follows:

1. Masking out the ESM-C feature corresponding to a single residue and recording the predicted probability *P*(*s*);
2. Inputting the complete ESM-C feature and retrieving the corresponding probability *P^*(*s*);
3. Computing the contribution score as *P^*(*s*) − *P*(*s*).

Based on this score, residues with the 10 highest values of *P^*(*s*) − *P*(*s*) were selected as the top-10 positively contributing residues. We further calculated the top-10 hit rate as follows:

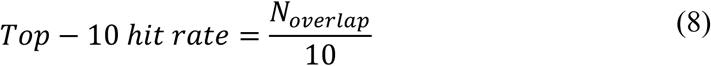

where *N_overlap_* denotes the number of residues among the top-10 positively contributing residues that overlap with annotated sites from UniProt.

For reaction-side interpretability, we extracted the local-global attention weights from the first layer and further averaged them over all enzyme residues to obtain an overall attention score for each reaction token. The scores were normalized and visualized as an attention map.

### Validation on external test sets

We next evaluated whether the locked model generalized beyond the development data by applying it, without further tuning, to independent external test sets. The first external test set, Enzyme-405 [62], comprises 961 positive enzyme-reaction pairs across 295 reactions and 405 enzymes, together with 14,960 negative pairs generated from Rhea snapshot (12 July 2023) [14]. For each reaction, the dataset provides a large candidate enzyme pool, closely matching the setting of enzyme screening. Following previous work [62], we evaluated the suitability of MAERM’s binary-classification probabilities for enzyme screening. We employed enrichment factor (EF), top-k success rate (SR) and top-n discounted cumulative gain (DCG), which measure positive-enzyme enrichment, retrieval success and ranking performance, respectively. The three metrics are defined as follows:

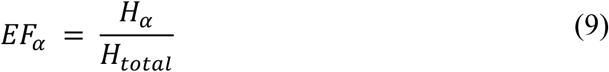

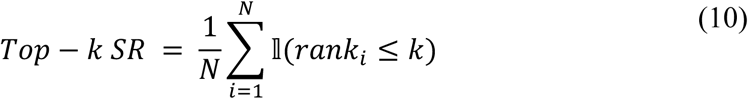

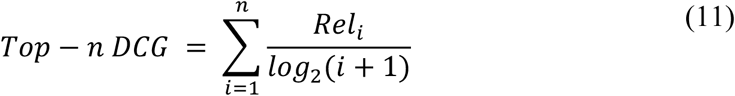

Here, *H_α_* and *H_total_* denote the hit rates within the top *α* fraction and the full dataset, respectively. *N* is the number of test cases, *rank_i_* is the rank of the correct prediction for the *i^th^* case, l is an indicator function, and *Rel*i is the ground-truth label (0 or 1) of the candidate at position *i*. All rankings are based on the probabilities predicted by MAERM. For top-k SR, we used k = 1, 3, 5, and 10; for top-n DCG, we used n = 5 and 10.

The datasets above still lack experimentally grounded negative examples, so we curated 547 enzyme-reaction pairs with reported conversion values from four studies [63–66]. The resulting dataset, BioCat-547, mainly comprises imine reductase (IRED)-and transaminase-catalyzed cases. We retained cases with conversions measured over 18-24 h and defined those with conversions below 30% as negative, yielding 193 positive and 354 negative pairs. For this imbalanced dataset, we evaluated performance using balanced accuracy, AUROC, AUPR, MCC and macro-F1 via Scikit-learn; metric definitions are provided in **Supporting Information Part 4**.

### Integrating ProteinMPNN for Enzyme Design and Screening

To investigate the ability of MAERM combined with ProteinMPNN [37] in enzyme design and screening, we selected *Candida parapsilosis* carbonyl reductase (CpRCR) as the template. The crystal structure of wild-type CpRCR (PDB ID: 3WLE) was used as the basis [67], and 27 mutation hotspots identified by full-sequence evolutionary analysis [68] were set as designable positions for ProteinMPNN. We selected the reduction reactions of the aryl ketone substrate A2 and the heterocyclic ketone substrate H4 from the literature as investigated reactions [68]. We prioritized these reactions because the previously reported active M3 variant (I51L/Y61F/D147E) exhibited only marginal enhancements in *k_cat_* relative to the wild type. In fact, these enhancements are among the lowest for analogous substrates, indicating significant potential for further optimization. For each investigated reaction, we evaluated designed enzymes using MAERM, and the top-ranking enzyme was selected based on the predicted probability.

Molecular dynamics (MD) simulations were conducted on the selected enzymes and the M3 variant with OpenMM 8.1 and AmberTools 24 [69, 70]. After equilibration, each system was subjected to a 200 ns production MD simulation. Active prereactive-state (PRS) conformations were calculated from the final 100 ns of each trajectory by mechanism-based geometric criteria [71], with a Bürgi-Dunitz attack angle of 107 ± 10° and a donor-acceptor distance ≤ 3 Å. Finally, we calculated the RTMScore [72] between the selected enzymes, the M3 variant, and their corresponding substrates, where a higher RTMScore indicates stronger predicted protein-ligand binding affinity. Details of the integration with ProteinMPNN and the MD simulation protocols are provided in **Supporting Information Part 5**.

## 3. Results

### Data analysis

We performed an in-depth analysis of MAERM-DB (Figure 3), which comprises 5,046 non-redundant reactions and 127,127 enzymes derived from both positive and negative samples. Examination of data sources reveals a significant imbalance, with most entries originating from ESP-ER (Figure 3a). Notably, the RDB dataset shares few enzymes or reactions with the other two sources (i.e., ESP-ER and M-CSA), reflecting the uniqueness of chemoenzymatic catalysis. Further analysis of reaction SMILES and enzyme sequence length distributions (Figure 3b, d) reveals a clear pattern. The RDB dataset contains shorter SMILES and enzyme sequences compared with ESP-ER and M-CSA. In contrast, no significant differences are observed in SMILES length between positive and negative samples (Figure 3c). Similarly, no differences are observed in enzyme sequence length across varying identity levels within the test set (Figure 3e). These results underscore a fundamental challenge in enzyme-reaction matching, as models cannot rely on superficial features such as sequence length for memorization. Instead, reliable prediction requires discerning intrinsic differences between positive and negative pairs.

**Figure 3.**
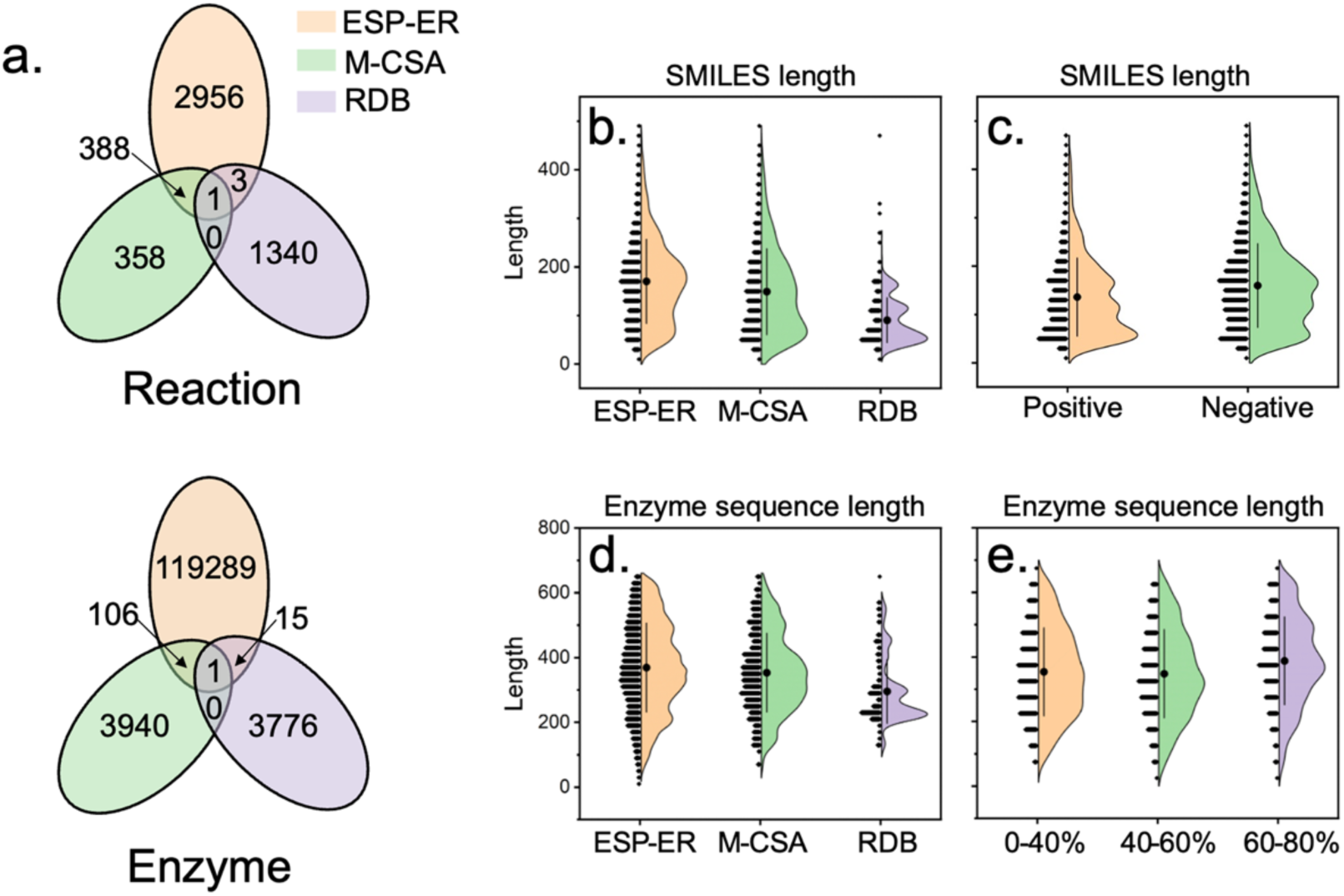
Analysis of MAERM-DB. (a) The distribution of non-redundant reactions and enzymes in MAERM-DB. (b) SMILES length for data from ESP-ER, M-CSA, and RDB datasets. (c) SMILES length for positive and negative samples. (d) Enzyme sequence length for data from ESP-ER, M-CSA, and RDB datasets. (e) Enzyme sequence length across different identity levels.

TMAP dimensionality reduction was applied to visualize the distribution of MAERM-DB (Figure 4a-d). In Figure 4a-c, the resulting trees are colored by data source (ESP-ER, M-CSA, and RDB), where neighboring nodes represent reactions, enzymes, or enzyme-reaction pairs with high similarity. Obviously, entries from RDB form well-separated clusters and occupy distinct regions. This pattern highlights the strong uniqueness of the RDB dataset, consistent with the results shown in Figure 3. Finally, Figure 4d is colored by positive and negative pairs. Representations corresponding to positive and negative pairs are highly intermixed, reflecting an overlapping distribution. This is expected because the negative reaction is defined as the most similar, non-identical reaction to a positive one, resulting in a median similarity of 0.983 between them (Figure S1). This overlapping distribution highlights the complexity of the prediction task and suggests that intrinsic differences between positive and negative pairs should be explored.

**Figure 4.**
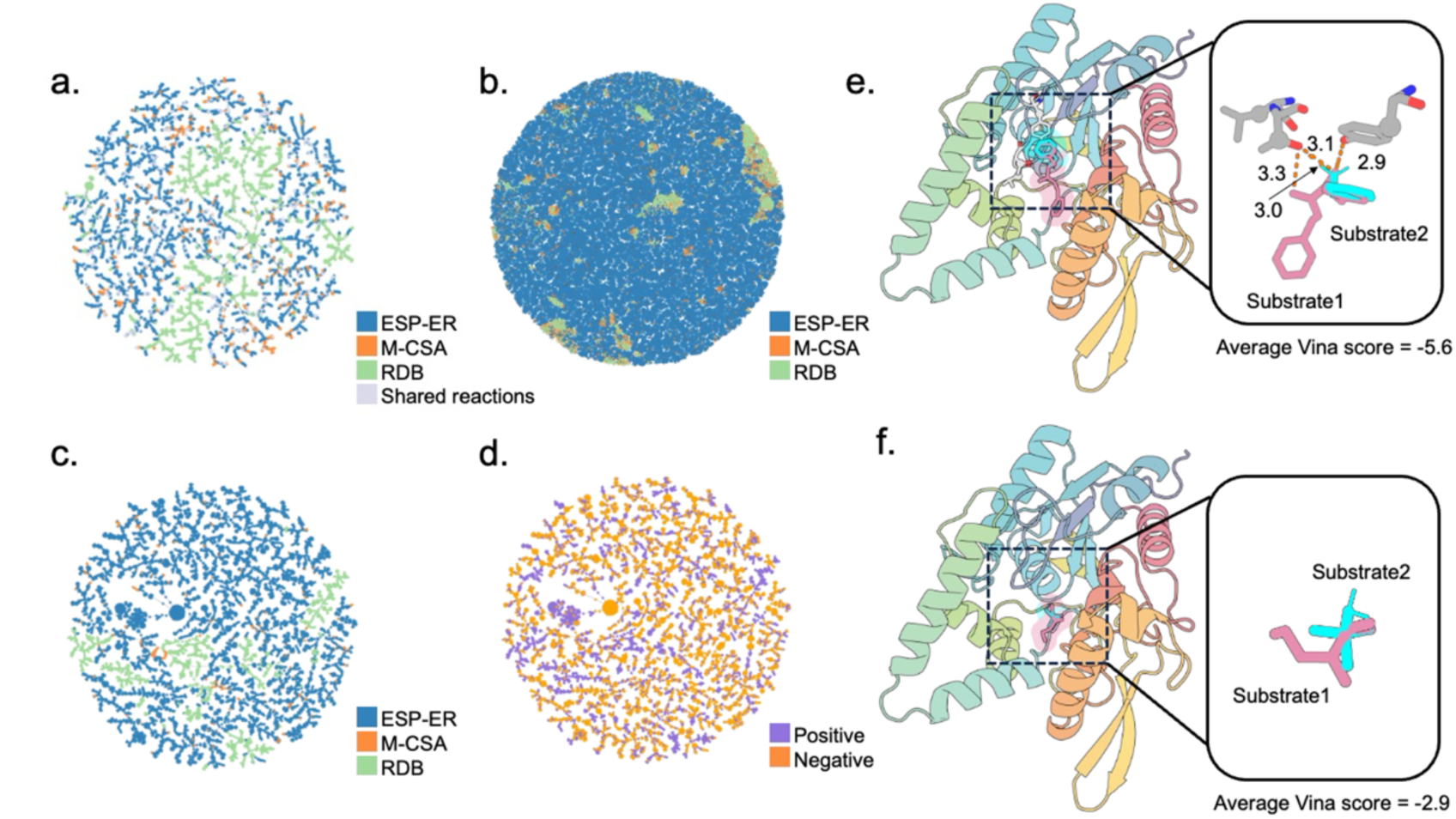
TMAP reduction and docking analysis. (a-b) TMAP visualization of reactions (rxnfp) and enzymes (ESM-C), color-coded by data source. (c-d) Concatenated enzyme-reaction features colored by data source and positive/negative label, respectively. To improve readability, 5,000 pairs were independently sampled from each of the positive and negative sets. (e-f) Docking of enzyme 1HQD with (e) positive and (f) negative substrates, showing polar interactions (< 3.5 Å) exclusively in the positive sample.

We evaluated our negative pair generation by calculating the average Vina scores for all enzymes and corresponding substrates in the RDB dataset. As shown in Figure S2, positive samples consistently exhibit lower Vina scores, indicating a stronger affinity between the substrates and enzymes. To illustrate these differences at the structural level, we visualized a representative enzyme (PDB ID: 1HQD) docked with positive and negative substrates (Figure 4e-f). Positive substrates form more polar interactions compared with negative counterparts, confirming their higher binding affinity. Overall, these results support the rationale of our negative pair generation, showing that the pairs remain chemically akin to positives yet display reduced catalytic potential.

### Results of enzyme-reaction matching prediction

We evaluated MAERM on MAERM-DB using a suite of benchmark experiments designed to quantify its ability to predict enzyme-reaction matching. On the overall test set, MAERM achieves the strongest performance among all evaluated methods, with an AUC of 0.999 and an F1-score of 0.984 (Table S5). Performance on enzymes with low identity to the training set (Figure 5a-c) provides a more stringent evaluation, as it assesses whether MAERM’s performance reflects learned biocatalytic principles rather than rote memorization. In the most stringent setting, where test enzymes share only 0-40% sequence identity with the training set (Table 2), MAERM consistently outperforms all baselines, surpassing MAARDTI by 3.7-7.3% and MCANet by 3.6-7.7% across five metrics. This ability is critical for real-world applications involving remote homologues, many of which preserve only functional catalytic centers and share less than 30% sequence identity [73].

**Figure 5.**
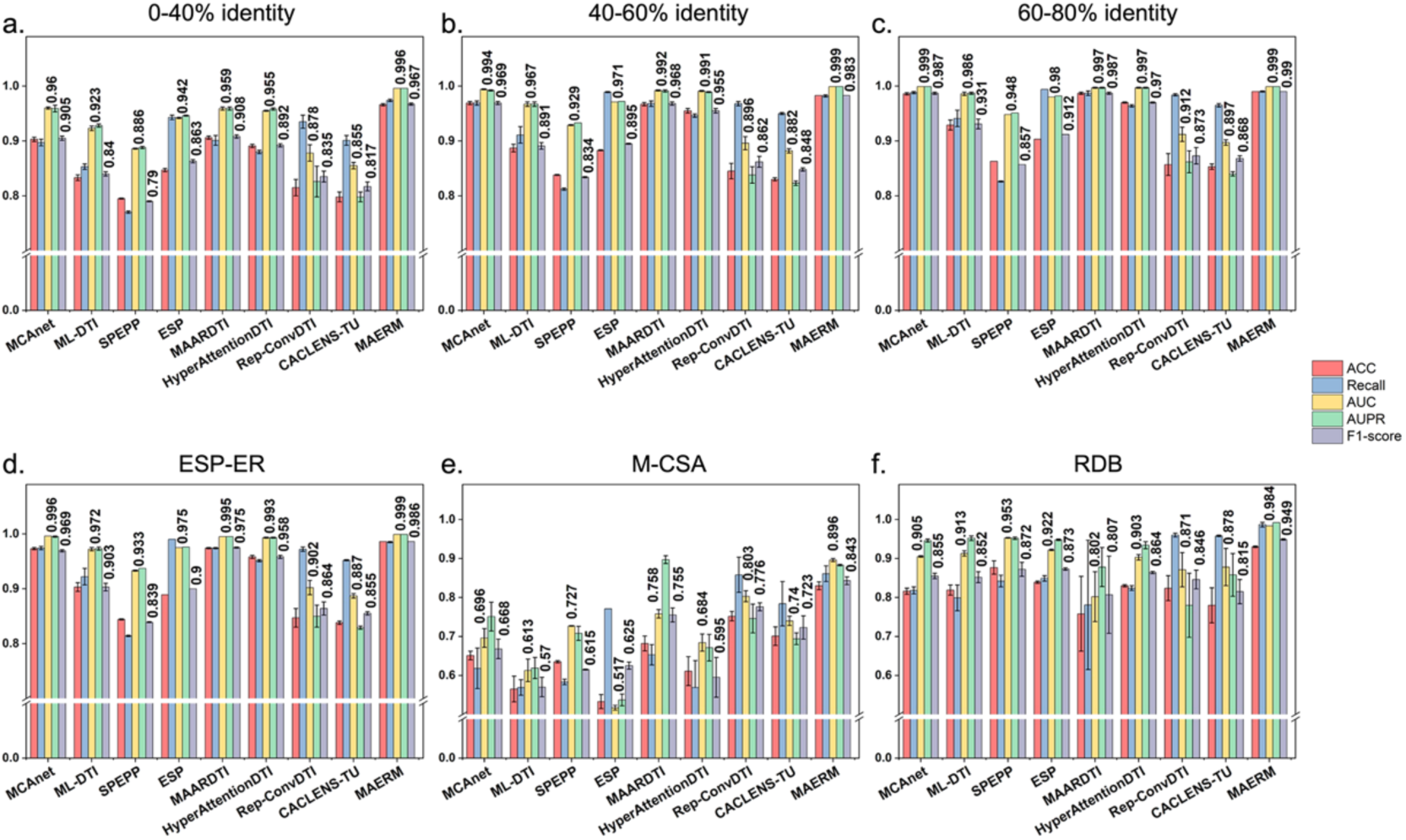
In-depth performance comparison of different models on the MAERM-DB dataset. (a-c) Model performance on test subsets stratified by enzyme sequence identity (i.e., 0-40%, 40-60%, and 60-80%). (d-f) Overall performance on test subsets derived from different data sources (i.e., ESP-ER, M-CSA, and RDB).

**Table 2.**
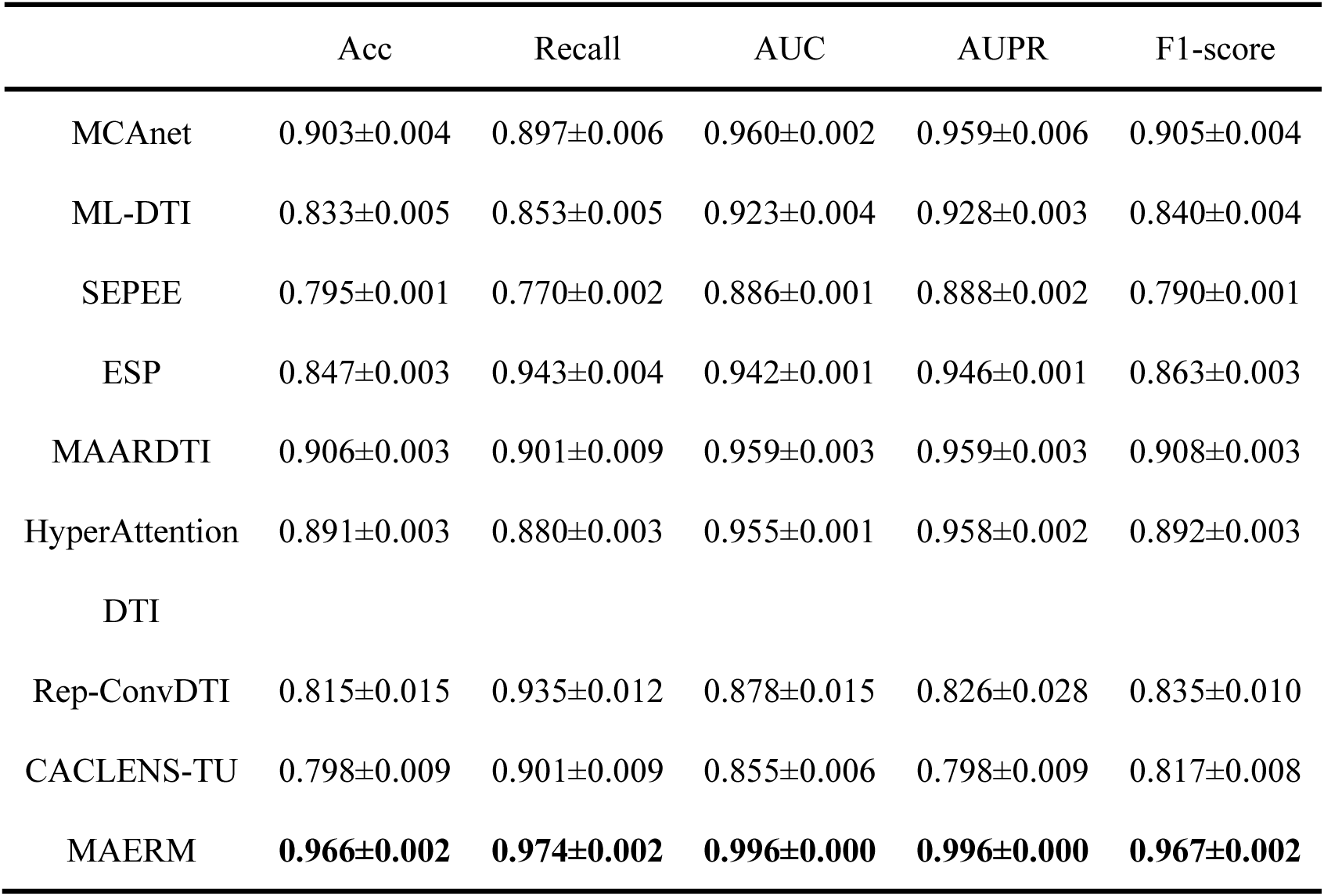
Performance comparison of different models on the MAERM-DB dataset with 0-40% sequence identity.

We further evaluated performance on test subsets from different data sources (ESP-ER, M-CSA, and RDB). As shown in Figure 5d-f and Table S6, MAERM consistently outperforms other models on the M-CSA and RDB subsets, surpassing the second-best model by 6.7% and 7.6% in F1-score, respectively. Focusing on the top-performing models in Table S5 (MAERM, MAARDTI, and MCANet), we compared their receiver operating characteristic (ROC) and precision-recall (PR) curves across these subsets. We selected the best run of these three models, as determined by the highest F1-score. As shown in Figure 6, MAERM achieves the best ROC and PR curves across all evaluation settings, with both AUC and AUPR values exceeding 0.88. In contrast, MAARDTI and MCANet show pronounced performance degradation on the M-CSA and RDB subsets. Together, these results show that MAERM performs well on both expert-curated data (M-CSA) and chemoenzymatic (RDB) data, supporting its generalizability across diverse application scenarios.

**Figure 6.**
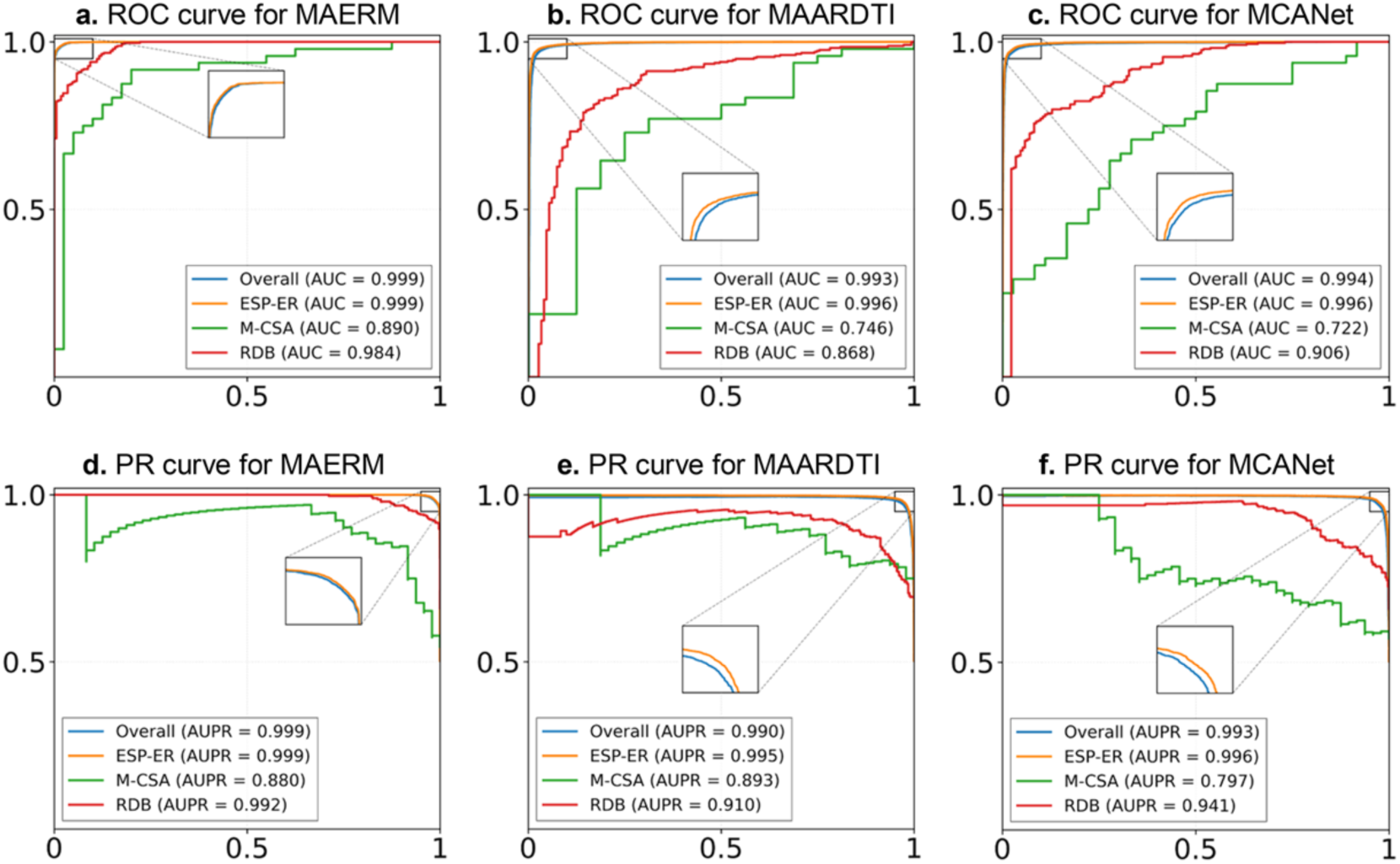
Comparative ROC and PR curve analysis across three data sources. (a-c) ROC curves for MAERM, MAARDTI, and MCANet across the three data sources. (d-f) PR curves for the same models and data sources.

### Results of ablation studies

We performed ablation studies to evaluate the contribution of individual components to enzyme-reaction matching performance (Table 3). Within the enzyme encoder, removing the geometry-sequence graph results in a 4.2% decrease in F1-score, indicating that geometry information helps capture spatial constraints beyond sequence-level representations. Our results also support the effectiveness of ESM-C features, as either excluding them (i.e., MAERM-w/o Seq) or replacing them with residue-level one-hot embeddings (i.e., MAERM-Residue) results in a notable decrease in performance. Within the reaction encoder, the effectiveness of the annealing strategy is observed. Specifically, consistently using information from reaction SMILES throughout training (i.e., MAERM-SMILES) or from the reaction center (i.e., MAERM-Center) reduces the F1-score by 3.8% and 0.7%, respectively. Finally, an ablation study on the reaction center radius shows that MAERM with the default radius (i.e., set to 1) achieves the best performance, outperforming both MAERM-R0 and MAERM-R2. This observation highlights a trade-off in defining the reaction center, with smaller radii potentially missing relevant side-chain context and larger radii introducing unrelated chemical groups and noise.

**Table 3.**
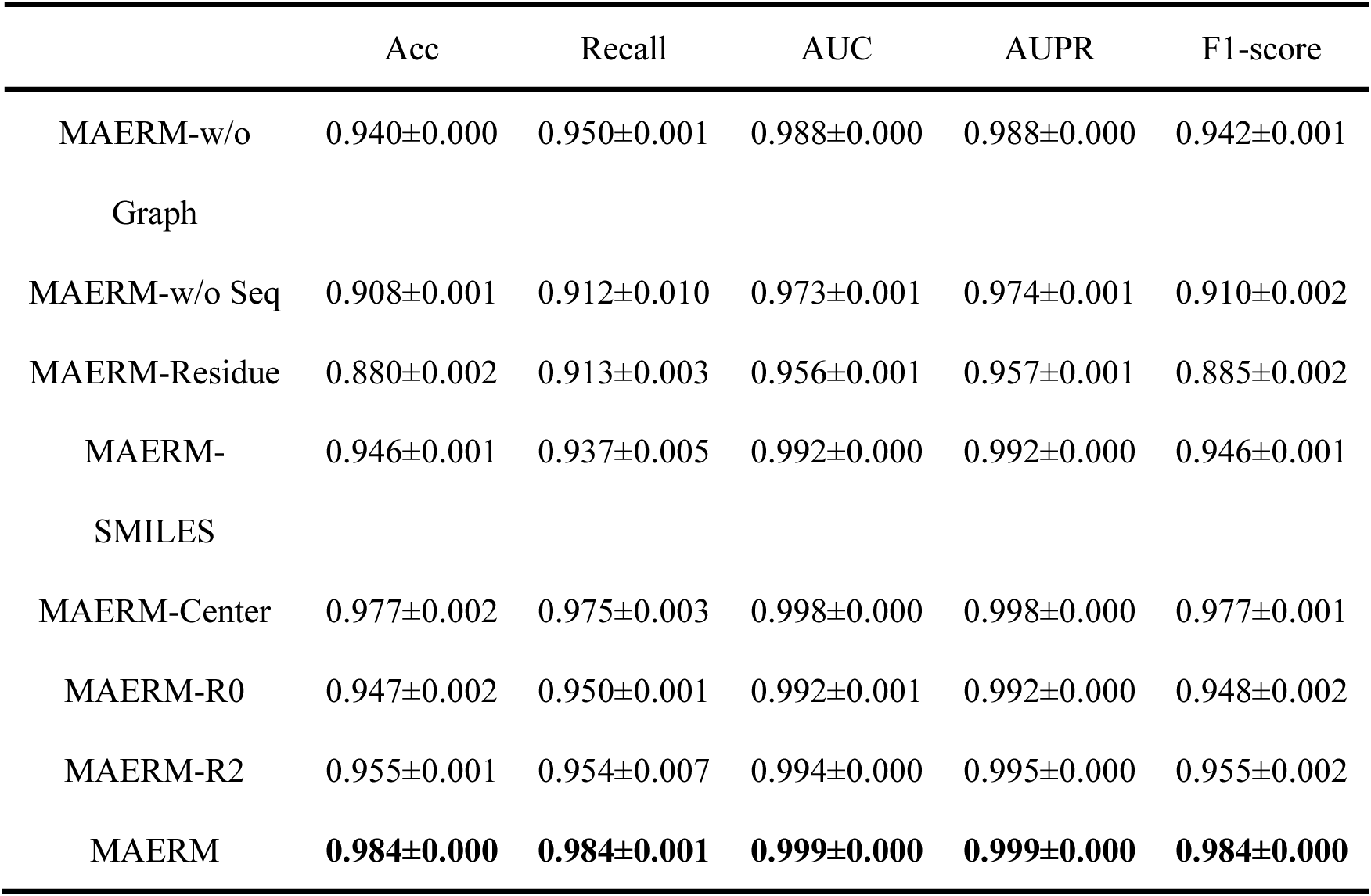
Ablation study of MAERM on the MAERM-DB dataset.

### Interpretability of MAERM

We analyzed the interpretability of the MAERM model using the best-performing run, as determined by the highest F1-score. To quantify residue contributions, we perturbed individual residues and measured the resulting change in enzyme-reaction matching probability. A larger drop in probability indicates a more important residue, whereas smaller or negligible changes suggest limited contribution. Figure 7 presents the results of perturbation-based interpretability analysis across representative enzyme-reaction pairs. Across these examples, the overlap between the top-10 positively contributing residues and annotated sites is consistently high, with hit rates ranging from 40% to 70%. In Q42972 (Figure 7b), for instance, the top-10 positively contributing residues include Arg124, Arg130, Asn162, Arg196, and Met271. Notably, UniProt annotates residues 124, 130, 162, and 196 as binding sites associated with the substrate, all of which are recovered. These results show that, even under challenging conditions lacking cofactor information, the model remains capable of capturing key enzymatic residues, underscoring its strong interpretability.

**Figure 7.**
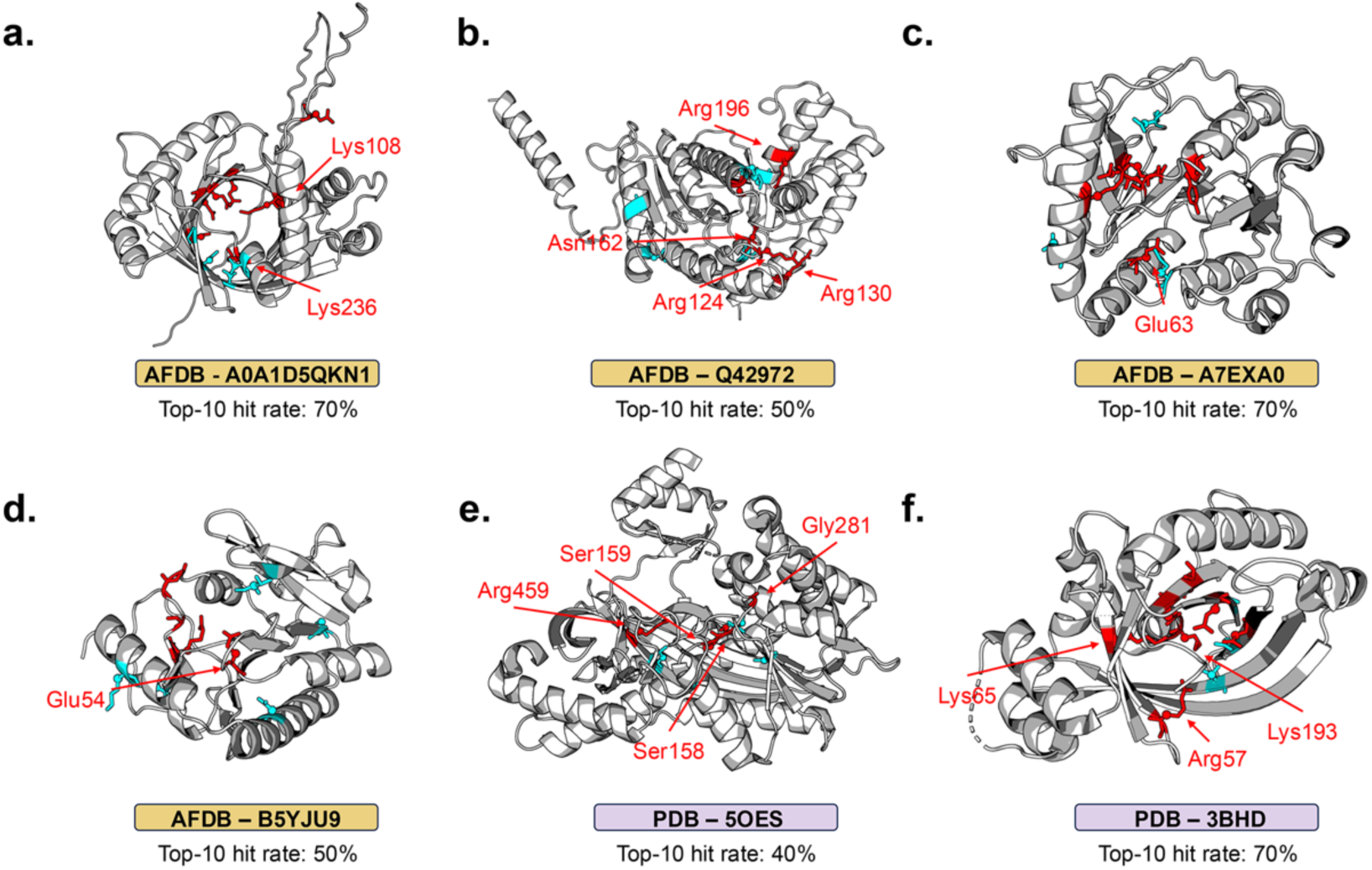
Perturbation-based interpretability analysis. Representative examples include: (a-d) A0A1D5QKN1, Q42972, A7EXA0, and B5YJU9 from AFDB; (e-f) 5OES and 3BHD from PDB. Top-10 positively contributing residues are shown in stick representation and colored red when they coincide with annotated UniProt sites, cyan when they do not. Annotated sites associated with the binding of substrate are labeled separately. Top-10 hit rate denotes the proportion of annotated sites among the top-10 positively contributing residues, as calculated in Equation 8.

For reaction-side interpretability, we examined the corresponding reaction attention in Figure S3-S5. In diphosphonic acid hydrolysis (Figure S3), the model predominantly attends to the phosphate bond, whereas in the transamination reaction (Figure S5), attention is focused on the amino group. Notably, beyond these reactive groups, the model also assigns substantial attention to the stereochemical symbols “@” or “@@”, suggesting that it captures stereochemical information associated with enzyme catalysis. Together, these observations provide deeper insight into how the model understands enzymatic catalysis.

### Results on the external test set

Figure 8 and Table S7 present data analysis and prediction results on the two external test sets, Enzyme-405 and BioCat-547. For data analysis, enzymes in BioCat-547 show lower sequence identity to the training set than those in Enzyme-405 (Figure 8a-b). BioCat-547 mainly contains chemoenzymes derived from multi-mutant engineering [63–66], most of which have not been recorded in existing datasets. These enzymes still show a degree of internal sequence similarity. For example, six representative imine reductases (IREDs) in BioCat-547 share several conserved regions (Figure S6). Additionally, both test sets are highly imbalanced (Figure 8c), reflecting real-world enzyme catalysis scenarios in which active enzymes are uncommon [64].

**Figure 8.**
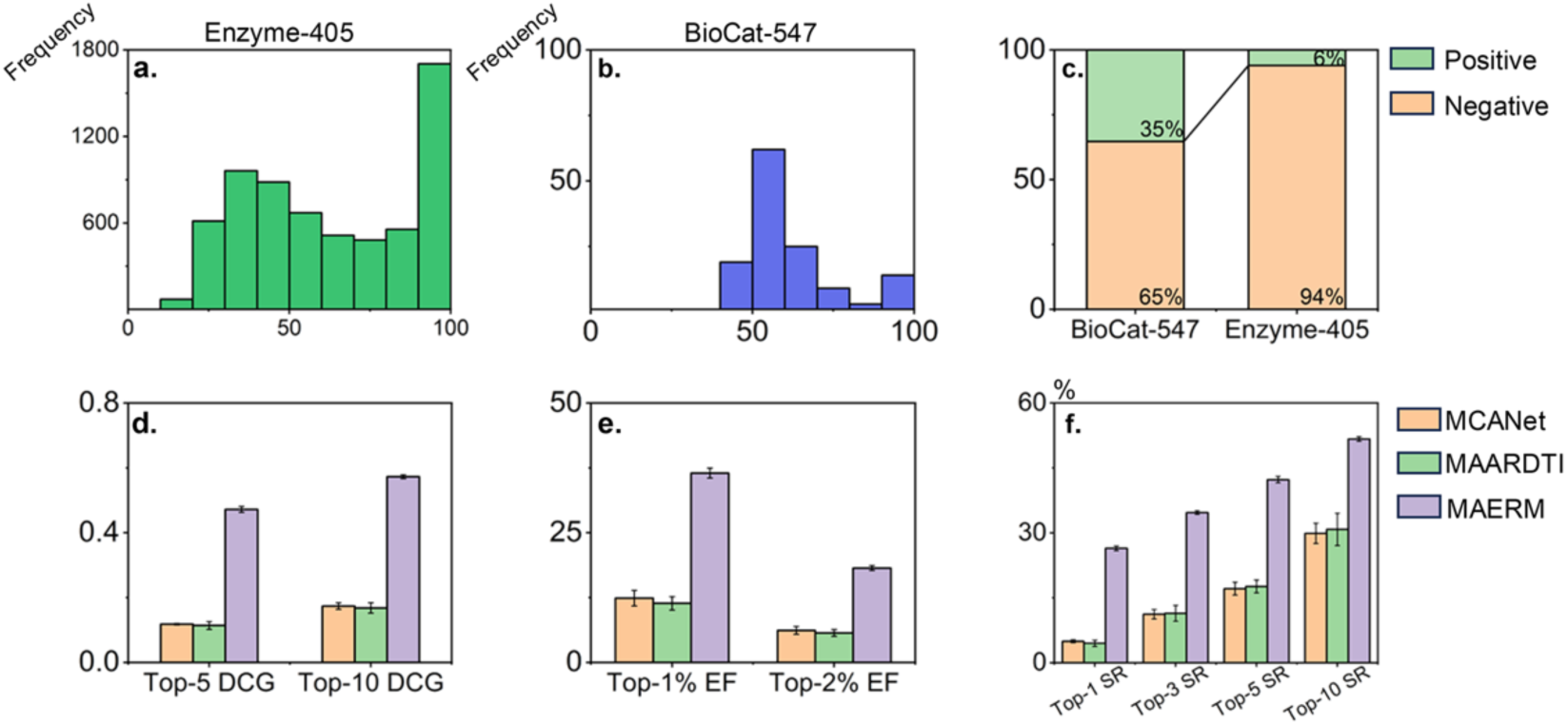
Data analysis and prediction results on the external test sets. (a-b) Sequence identity between enzymes in the training set and those in the Enzyme-405 and BioCat-547 datasets, respectively, as calculated by MMseqs2 [74]. (c) Distribution of positive and negative samples in the Enzyme-405 and BioCat-547 datasets. (d-f) Performance of MAERM and baseline models on Enzyme-405, including top-n DCG, EF, and top-k SR.

On the Enzyme-405 dataset (Figure 8d-f), MAERM achieves an average top-10 DCG of 0.573, a top-2% EF of 18.2, and a top-10 SR of 51.7%, substantially outperforming both baseline models. These results demonstrate that MAERM enriches active enzymes among top-ranked candidates, highlighting the utility of its predicted probabilities for guiding downstream enzyme screening. On BioCat-547 (Table S7), MAERM also achieves the best performance across all metrics, with a balanced accuracy of 0.697. This value exceeds that of MAARDTI by 0.18, supporting MAERM’s generalizability to chemoenzymatic catalysis.

To further assess the performance of MAERM, we have selected five cases from BioCat-547 for visualization. As shown in Figure 9, our model successfully identifies three IRED-catalyzed cases and two transaminase-catalyzed cases. Moreover, the predicted probabilities are generally consistent with real conversion rates. These results further indicate that MAERM maintains reliable predictive performance on experimentally validated samples.

**Figure 9.**
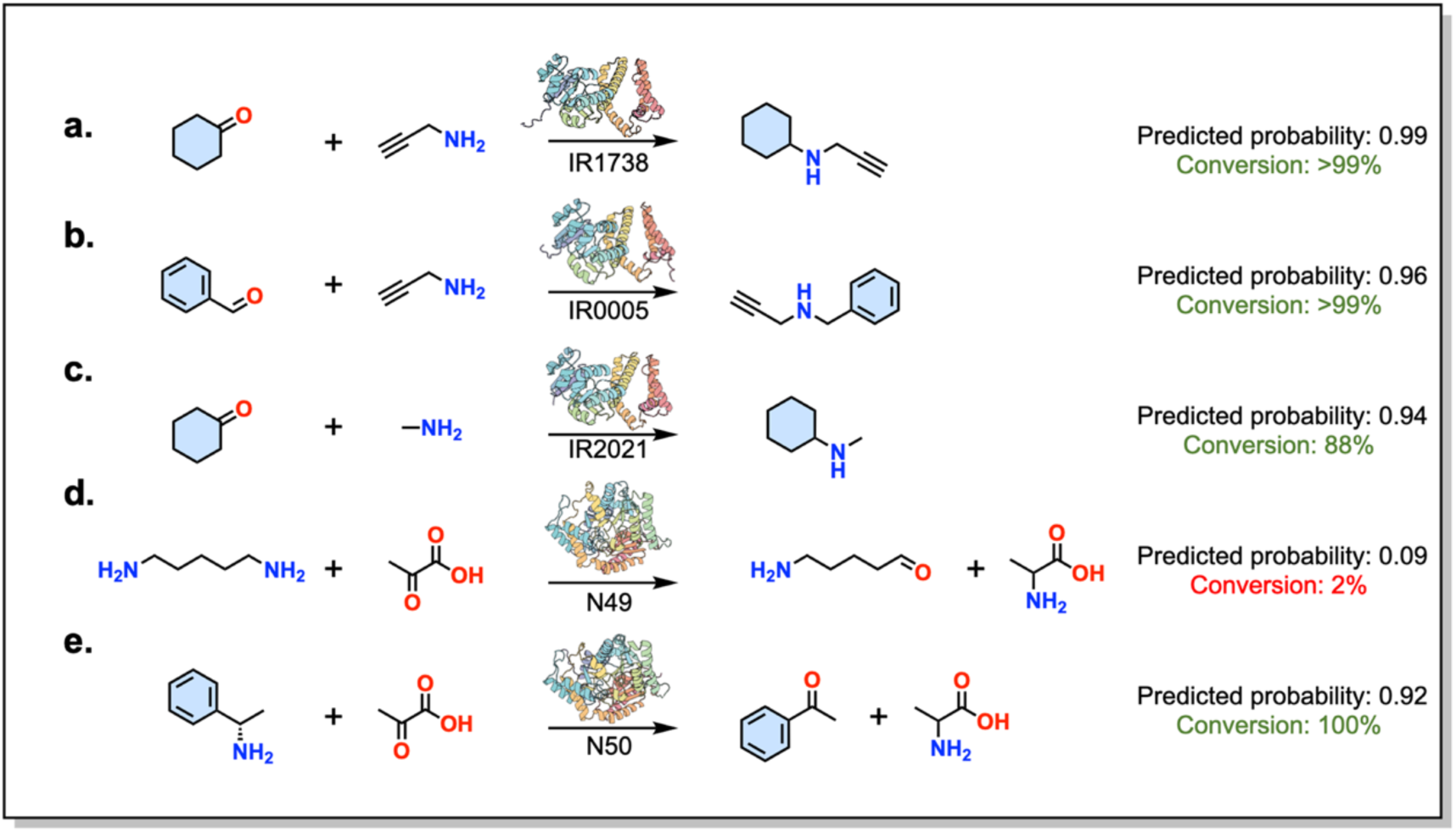
Five representative cases from the BioCat-547 dataset. (a-c) show three IRED-catalyzed cases, whereas (d) and (e) show two transaminase-catalyzed cases. The corresponding enzyme names are obtained from the literature [64, 66].

### Integrating ProteinMPNN for Enzyme Design and Screening

Integrating MAERM with protein design tools such as ProteinMPNN can help accelerate the design-build-test-learn cycle in enzyme engineering. Using wild-type CpRCR (PDB ID: 3WLE) as the template, we restricted sequence redesign to 27 mutational hotspots and generated enzyme variants with ProteinMPNN (Figure S7). Among these variants, MAERM identifies the top-ranking enzymes for two carbonyl reduction reactions. The reactions are shown in Figure 10a-b, and the corresponding enzymes in Figure 10c-d. Notably, both selected enzymes harbor the Y61F and D147E mutations. It has been suggested that these mutations balance global protein fluctuations by redistributing conformational flexibility across distinct regions [68].

**Figure 10.**
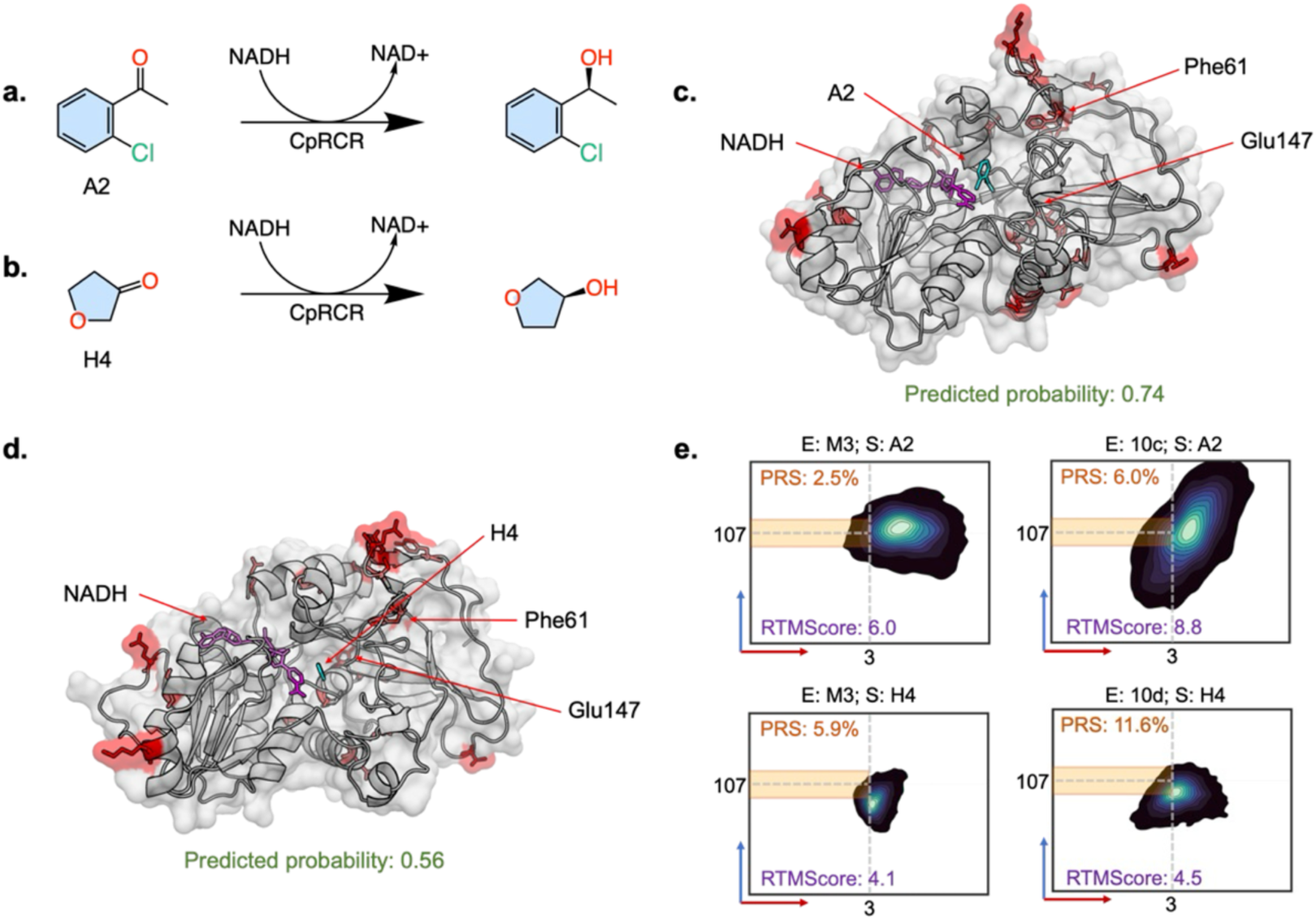
Enzyme design and validation of carbonyl reductase. (a-b) Reduction reactions of the aryl ketone substrate A2 and the heterocyclic ketone substrate H4, respectively, reported in the literature [68]. (c-d) Complex structures of A2 and H4 with the corresponding top-ranking enzymes. Residues redesigned by ProteinMPNN are shown in red. (e) MD simulations of the enzymes shown in 10c, 10d, and M3 variant (I51L/Y61F/D147E). The x- and y-axes represent the donor-acceptor distance and the Bürgi-Dunitz attack angle, respectively. The yellow region denotes PRS conformations. E and S denote the enzyme and its corresponding substrate, respectively.

We conducted molecular dynamics (MD) simulations for top-ranking enzymes and the literature-reported highly active M3 variant (I51L/Y61F/D147E). Kernel density analysis (Figure 10e) shows that top-ranking enzymes exhibit higher PRS populations of 6.0% and 11.6% for the two reduction reactions, respectively, whereas the M3 variant shows only 2.5% and 5.9%. These results indicate that top-ranking enzymes more densely sample catalytically competent conformations, making them more likely to catalyze the corresponding reactions. RTMScore analysis provides complementary support for this trend, revealing higher substrate affinity for top-ranking enzymes. Specifically, these enzymes show higher binding scores toward A2 and H4 than the M3 variant, with RTMScores of 8.8 and 4.5, respectively. Given that M3 is obtained through multiple rounds of greedy directed evolution and experimental validation, the combination of ProteinMPNN and MAERM provides a more efficient strategy for exploring combinatorial sequence space and capturing additive or synergistic effects among designed sites.

## 4. Conclusion

This study introduces MAERM, a novel mixed-attention model designed to predict enzyme-reaction matching relationships. By coupling multimodal enzyme information (i.e., geometry-sequence graphs and ESM-C features) with fine-grained reaction representations (i.e., local reactive regions and global reaction contexts), our model achieves marked improvements in predictive performance, especially on enzymes with low identity to the training set. Additionally, the model provides a degree of interpretability by capturing specific residues aligned with annotated binding sites. Evaluations on external test sets demonstrate that MAERM significantly outperforms baseline models across diverse scenarios, including enzyme screening and chemoenzymatic catalysis. Furthermore, by successfully integrating ProteinMPNN with MAERM, we demonstrate that MAERM can serve as an efficient scoring module to guide enzyme design for two carbonyl reduction reactions.

Our research makes a significant contribution by curating a new dataset, MAERM-DB, which offers broad insights into data reliability and coverage. To fully leverage this robust dataset, our model integrates multimodal enzyme information with fine-grained reaction representations via a mixed local-global attention module. This enables the model to learn both the diverse enzyme syntax and the multi-level contexts of reactions. Compared with earlier studies [26–28], MAERM not only achieves higher accuracy and interpretability but also demonstrates broad compatibility, seamlessly integrating into diverse downstream scenarios.

The primary challenge in this research remains the risk of false negatives. Owing to the limited availability of well-curated negative samples, MAERM uses generated enzyme-reaction pairs as negatives for training, which may not capture the complete catalytic scope of an enzyme. Nevertheless, MAERM achieves highly competitive performance on BioCat-547, which contains experimentally validated negative samples, supporting its reliability and practical relevance. At the modeling level, some recent studies have incorporated computational enzyme annotations such as binding-pocket information [62], to emphasize functionally relevant structural features. However, the widespread gaps in enzyme functional annotations mean that relying solely on computational methods inevitably leads to uncertainty and added computational costs [75]. Our future work will explore alternative enzyme representations, with particular attention to the trade-off between diverse representations and computational cost.

In conclusion, the MAERM model proposed in this study offers a fresh perspective on enzyme-reaction matching prediction. We hope this model can reduce the experimental costs of measuring enzymes’ catalytic scope, advance enzyme design, and ultimately accelerate the design-build-test-learn cycle in enzyme engineering.

## Additional information

### Publisher’s Note

Springer Nature remains neutral with regard to jurisdictional claims in published maps and institutional affiliations.

### Supporting Information

Additional Information (figures, tables and methodological details) is available in Supporting information. **Part 1.** The details of enzyme encoder; **Part 2.** The details of local-global attention module; **Part 3.** Baseline models used in this study; **Part 4.** The definitions of evaluation metrics; **Part 5.** Details of enzyme design and screening; **Part 6.** Supplementary Tables; **Part 7.** Supplementary Figures.

### Data Availability

The datasets supporting this study are available in Zenodo at https://doi.org/10.5281/zenodo.20151006. The source code required to reproduce the results is available at GitHub (https://github.com/Tiantao2000/MAERM).

### Competing interests

The authors declare no competing financial or non-financial interests.

### Author Contributions

T. L. and S. Z. contributed to the conceptualization and writing of the manuscript. S. L. and X. Z. contributed to revising the manuscript and data curation. J. D. contributed to data analysis. H. L. reviewed the manuscript and provided valuable guidance. Shirley W. I. Siu supervised the study and ensured the overall quality of the work.

### Funding

This project was funded by Macao Polytechnic University with grant number RP/FCA-18/2025. The funders had no role in study design, data collection, and interpretation, or the decision to submit the work for publication. This project is part of the thesis work of T. L. with an internal reference number of s/c fca.1d25.b5b0.c.

## Acknowledgements

We would like to sincerely thank the authors of the resources used in this study for their valuable contributions in sharing datasets with the scientific community.

## Author information

Faculty of Applied Sciences, Macao Polytechnic University, 999078, Macau SAR, China

Tiantao Liu, Silong Zhai, Shaolong Lin, Xinke Zhan, Junwen Deng, Huanxiang Liu, Shirley W. I. Siu

College of Pharmaceutical Sciences, Zhejiang University, Hangzhou 310058, Zhejiang, China

Silong Zhai

## Corresponding authors

Correspondence to <u>Shirley W. I. Siu</u>.

## Notes

https://github.com/Tiantao2000/MAERM

https://doi.org/10.5281/zenodo.20151006

### Competing Interest Statement

The authors have declared no competing interest.

