## Supporting Information for "MAERM: Predicting Enzyme-Reaction Matching Relationships with a Mixed-Attention Model"

---

Tiantao Liu[1], Silong Zhai[1,2], Shaolong Lin[1], Xinke Zhan[1], Junwen Deng[1], Huanxiang Liu[1], Shirley W. I. Siu[1,*]

[1]Faculty of Applied Sciences, Macao Polytechnic University, 999078, Macau SAR, China
[2]College of Pharmaceutical Sciences, Zhejiang University, Hangzhou 310058, Zhejiang, China

---

**Part 1. The details of enzyme encoder**

Enzyme encoder generates enzyme embeddings by integrating the feature of ESM-C [1] and geometry-sequence graph [2] through a cross-attention module.

In the enzyme encoder, we used ESM-C (600M, version 2024-12), which consists of 36 layers and 18 attention heads. In the geometry-sequence graph branch, the geometric radius was set to [4.0, 10.0], the sequential kernel size to 21, the bottleneck width to 32, and the channel dimensions to [256, 512]. Features from ESM-C and the geometry-sequence graph were fused through a cross-attention module with 8 heads and normalized to produce representations of 256 dimensions.

**Part 2. The details of local-global attention module**

The local-global attention module comprises two layers with eight attention heads, including four local and four global heads. The values of d_model, d_ff, and the dropout rate were set to 256, 1024, and 0.2, respectively. The output representations were normalized by layer normalization and aggregated by mean pooling over all nodes. The pooled 256-dimensional representation was then fed into a multi-layer perceptron (MLP) classifier for binary classification. Cross-entropy loss was used between the predicted probabilities and labels. Model training was conducted using stochastic gradient descent (SGD) with an initial learning rate of 1e-3, momentum of 0.9, and weight decay of 5e-4. The learning rate was decayed by a factor of 0.5 at epochs 20, 40, 60, and 80 using a LambdaLR scheduler. The model was trained for 100 epochs with a batchsize of 32 using NVIDIA A800 GPU and PyTorch 2.4.1+cu124, each experiment was repeated three times.

**Part 3. Baseline models used in this study**

We selected SPEPP [3], CACLENS [4], ESP [5], MCANet [6], ML-DTI [7], MAARDTI [8], and Rep-ConvDTI [9] as our baseline models. The default hyperparameters and code can be found at:

https://zenodo.org/records/8210150, https://github.com/XilongYi/CACLENS, https://github.com/AlexanderKroll/ESP, https://github.com/MrZQAQ/MCANet, https://github.com/guaguabujianle/ML-DTI, https://github.com/TorchZhan/MAARDTI, and https://github.com/DMP321/Rep-ConvDTI.

All baseline models were retrained three times on the MAERM-DB dataset using a single NVIDIA A800 GPU.

### Part 4. The definitions of evaluation metrics

We evaluated model performance using six standard metrics for MAERM-DB: accuracy (Acc), recall, precision, F1-score, area under the receiver operating characteristic curve (AUROC), and area under the precision-recall curve (AUPR). All metrics were calculated using Scikit-learn [10]. The definitions are as follows:

$$Acc = \frac{TP + TN}{TP + TN + FP + FN} \tag{1}$$

$$Precision = \frac{TP}{TP + FP} \tag{2}$$

$$Recall = \frac{TP}{TP + FN} \tag{3}$$

$$\mathrm{F1\_score} = 2 \times \frac{Precision \times Recall}{Precision + Recall} \tag{4}$$

where $TP$, $TN$, $FP$, and $FN$ denote true positives, true negatives, false positives, and false negatives, respectively. AUROC summarizes the trade-off between the true positive rate and false positive rate across different decision thresholds, whereas AUPR summarizes the trade-off between precision and recall across thresholds.

For the imbalanced BioCat-547 dataset, in addition to AUROC and AUPR, we further introduced balanced accuracy, MCC, and macro-F1 for evaluation. Their definitions are as follows:

$$\text{Balanced Accuracy} = \frac{1}{2}\left(\frac{TP}{TP+FN} + \frac{TN}{TN+FP}\right) \tag{5}$$

$$MCC = \frac{TP \times TN - FP \times FN}{\sqrt{(TP+FP)(TP+FN)(TN+FP)(TN+FN)}} \tag{6}$$

$$Macro\_F1 = \frac{F1_{positive} + F1_{negative}}{2} \tag{7}$$

**Part 5. Details of enzyme design and screening**

Based on previous studies [11], we selected the following 27 mutation hotspots in wild-type CpRCR (PDB ID: 3WLE): 13N, 14K, 15Q, 19K, 22N, 51I, 61Y, 72A, 76D, 77D, 79I, 138S, 146A, 147D, 152S, 204K, 221T, 246V, 266N, 272G, 293D, 294D, 304V, 310S, 314K, 320I, and 327A. In ProteinMPNN, we utilized NUM_SEQ_PER_TARGET = 500000, BATCH_SIZE = 50, and SAMPLING_TEMP = 0.3, which yielded 229,078 nonredundant sequences. The structure of each sequence was then predicted using ESM3-small (version 2024-08) [12]. The reaction SMILES of the aryl ketone substrate A2 and the heterocyclic ketone substrate H4 reported in the literature [11] are shown below:

CC(=O)c1ccccc1Cl>>C[C@H](O)c1ccccc1Cl (Figure 10a);

O=C1CCOC1>>O[C@H]1CCOC1 (Figure 10b).

For these two reactions, MAERM was used to screen candidate enzymes, identifying the top-scoring variants as:

N13D/Q15K/K19Q/N22T/Y61F/I79T/S138V/A146P/D147E/S152C/T221E/V246I/G272I/D293E/D294E/K314E/I320L/A327E (Figure 10c);

and

N13S/Q15E/K19Q/N22Y/Y61F/I79T/S138V/A146F/D147E/S152C/T221E/V246I/G272K/D293E/V304I/K314I/I320L/A327E (Figure 10d).

Prior to molecular dynamics (MD) simulations, we first predicted the complex structure using AlphaFold3 [13]. After that, the protein was described with the AMBER ff14SB force field [14], whereas the substrate and Nicotinamide Adenine Dinucleotide Hydride (NADH) were parameterized using GAFF2 [15]. Each complex was solvated in TIP3P water [16], neutralized with $Na^+$ and $Cl^-$ ions, minimized, and equilibrated under NVT followed by stepwise NPT conditions. Long-range electrostatics were treated with particle mesh Ewald [17], and temperature was controlled using a Langevin thermostat [18]. After equilibration, each system was subjected to a 200 ns production simulation.

### Part 6. Supplementary Tables

**Table S1** Common co-enzymes and byproducts that are removed from the dataset

| Name | SMILES/SMARTS |
|---|---|
| Coenzyme A | O=C(NCC*)CCNC(=O)C(O)C(C)(C)COP(=O)(*)OP(=O)(*)OC*3O*(n2cnc1c(ncnc12)N)*(O)*3OP(=O)(*)* |
| Nicotinamide adenine dinucleotides | **1*(*)*(COP(*)(=O)OP(*)(=O)OC*2O*(*)*(*)*2*)O*1* |
| Nucleoside phosphates | **1*(*)*(O*1COP(*)(=O)O)[R] |
| Nucleoside phosphates isomers | *P(*)(=O)O*1*(*)*(*)O[*]1COP(*)(*)=O |
| Sulfonium betaines | **1*(*)*(O*1CS*)[R] |
| Flavines | **1**2**3*(**(=O)**3=O)*(*)*2**1* |
| Hemes | *~1~*~*~2~*~*~1~*~*~1~*~*~*(~*~*~3~*~*~*(~*~*~4~*~*~*(~*~2)~*~4)~*~3)~*~1 |
| Iron-sulfur cluster(s) | S1[Fe]S[Fe]1 |
| Phosphate trianion | O=P([O-])([O-])[O-] |
| Hydrogen phosphate | O=P([O-])([O-])O |

| | |
|---|---|
| dianion | |
| (2-hydroxyethyl) trimethylammonium | C[N+](C)(C)CCO |
| Ethanolamine | NCCO |
| Diphosphate | O=P([O-])([O-])OP(=O)([O-])[O-] |
| Hydrogen diphosphate trianion | O=P([O-])([O-])OP(=O)([O-])O |
| 2-Oxoglutarate dianion | O=C([O-])CCC(=O)C(=O)[O-] |
| Acetate ion | CC(=O)[O-] |
| Pyruvate | CC(=O)C(=O)[O-] |

**Table S2** Augmentation methods for M-CSA and RDB dataset

| | Definition | Proportion |
|---|---|---|
| Random Substitution | Randomly choose residues and replace them with other residues | 10% |
| Random Insertion | Insert residues into random positions | 5% |
| Random Deletion | Delete residues in random positions | 5% |
| Random Swap | Randomly choose two residues in the sequence and swap their positions | 5% |

**Table S3** The number of enzyme-reaction pairs retained after processing

| | MAERM-DB | ESP-ER | M-CSA | RDB |
|---|---|---|---|---|
| Original | 174172 | 169339 | 753 | 4080 |
| Positive Samples | 140128/14050/13598[a] | 105836/13615/13217 | 3952/49/48 | 30340/386/333 |
| Negative Samples | 122759/13722/13263 | 104327/13437/13040 | 3933/48/47 | 14499/237/176 |
| All Samples | 262887/27772/26 | 210163/27052/26 | 7885/97/95 | 44839/623/509 |

| | | |
|---|---|---|
| | 861 | 257 |

[a] The numbers from left to right are for the training set, validation set, and test set, respectively.

**Table S4** The number of enzyme-reaction pairs categorized by three levels of sequence identity in the test set

| | MAERM-DB | ESP-ER | M-CSA | RDB |
|---|---|---|---|---|
| 0-40% | 3252 | 3036 | 64 | 152 |
| 40-60% | 11117 | 10884 | 14 | 219 |
| 60-80% | 12492 | 12337 | 17 | 138 |

**Table S5** Performance comparison of different models on the MAERM-DB dataset.

| | Acc | Recall | AUC | AUPR | F1-score |
|---|---|---|---|---|---|
| **0-40%** | | | | | |
| MCAnet | 0.903±0.004 | 0.897±0.006 | 0.960±0.002 | 0.959±0.006 | 0.905±0.004 |
| ML-DTI | 0.833±0.005 | 0.853±0.005 | 0.923±0.004 | 0.928±0.003 | 0.840±0.004 |
| SEPEE | 0.795±0.001 | 0.770±0.002 | 0.886±0.001 | 0.888±0.002 | 0.790±0.001 |
| ESP | 0.847±0.003 | 0.943±0.004 | 0.942±0.001 | 0.946±0.001 | 0.863±0.003 |
| MAARDTI | 0.906±0.003 | 0.901±0.009 | 0.959±0.003 | 0.959±0.003 | 0.908±0.003 |
| HyperAttention DTI | 0.891±0.003 | 0.880±0.003 | 0.955±0.001 | 0.958±0.002 | 0.892±0.003 |
| Rep-ConvDTI | 0.815±0.015 | 0.935±0.012 | 0.878±0.015 | 0.826±0.028 | 0.835±0.010 |
| CACLENS-TU | 0.798±0.009 | 0.901±0.009 | 0.855±0.006 | 0.798±0.009 | 0.817±0.008 |
| MAERM | **0.966±0.002** | **0.974±0.002** | **0.996±0.000** | **0.996±0.000** | **0.967±0.002** |
| **40-60%** | | | | | |
| MCAnet | 0.969±0.003 | 0.969±0.004 | 0.994±0.001 | 0.992±0.001 | 0.969±0.003 |
| ML-DTI | 0.887±0.007 | 0.911±0.015 | 0.967±0.004 | 0.967±0.004 | 0.891±0.006 |
| SEPEE | 0.838±0.001 | 0.812±0.002 | 0.929±0.001 | 0.933±0.000 | 0.834±0.001 |

| | | | | | |
|---|---|---|---|---|---|
| ESP | 0.883±0.001 | **0.989±0.001** | 0.971±0.000 | 0.972±0.000 | 0.895±0.001 |
| MAARDTI | 0.967±0.003 | 0.968±0.005 | 0.992±0.001 | 0.991±0.002 | 0.968±0.003 |
| HyperAttention DTI | 0.955±0.004 | 0.946±0.003 | 0.991±0.001 | 0.989±0.001 | 0.955±0.004 |
| Rep-ConvDTI | 0.845±0.014 | 0.968±0.004 | 0.896±0.012 | 0.838±0.015 | 0.862±0.010 |
| CACLENS-TU | 0.830±0.003 | 0.950±0.002 | 0.882±0.004 | 0.823±0.004 | 0.848±0.003 |
| MAERM | **0.983±0.000** | 0.982±0.002 | **0.999±0.000** | **0.999±0.000** | **0.983±0.000** |
| **60-80%** | | | | | |
| MCAnet | 0.986±0.002 | 0.988±0.002 | 0.999±0.000 | 0.999±0.000 | 0.987±0.002 |
| ML-DTI | 0.929±0.009 | 0.941±0.015 | 0.986±0.003 | 0.987±0.002 | 0.931±0.009 |
| SEPEE | 0.863±0.000 | 0.826±0.001 | 0.948±0.000 | 0.951±0.000 | 0.857±0.000 |
| ESP | 0.903±0.000 | **0.994±0.000** | 0.980±0.000 | 0.982±0.000 | 0.912±0.000 |
| MAARDTI | 0.987±0.002 | 0.987±0.004 | 0.997±0.001 | 0.997±0.001 | 0.987±0.002 |
| HyperAttention DTI | 0.970±0.001 | 0.964±0.002 | 0.997±0.001 | 0.997±0.001 | 0.970±0.001 |
| Rep-ConvDTI | 0.857±0.020 | 0.984±0.002 | 0.912±0.013 | 0.862±0.020 | 0.873±0.015 |
| CACLENS-TU | 0.853±0.005 | 0.965±0.003 | 0.897±0.005 | 0.840±0.004 | 0.868±0.005 |
| MAERM | **0.990±0.000** | 0.990±0.001 | **0.999±0.000** | **0.999±0.000** | **0.990±0.000** |
| **Overall** | | | | | |
| MCAnet | 0.969±0.002 | 0.969±0.003 | 0.994±0.000 | 0.993±0.001 | 0.969±0.002 |
| ML-DTI | 0.900±0.008 | 0.918±0.013 | 0.972±0.003 | 0.973±0.003 | 0.903±0.007 |
| SEPEE | 0.844±0.000 | 0.813±0.001 | 0.933±0.001 | 0.936±0.000 | 0.839±0.000 |
| ESP | 0.888±0.000 | **0.986±0.000** | 0.973±0.000 | 0.975±0.000 | 0.899±0.000 |
| MAARDTI | 0.968±0.003 | 0.968±0.005 | 0.992±0.001 | 0.990±0.001 | 0.969±0.003 |
| HyperAttention DTI | 0.954±0.003 | 0.946±0.001 | 0.991±0.001 | 0.990±0.001 | 0.954±0.003 |
| Rep-ConvDTI | 0.846±0.017 | 0.971±0.004 | 0.901±0.013 | 0.848±0.019 | 0.864±0.012 |
| CACLENS-TU | 0.836±0.004 | 0.951±0.001 | 0.886±0.004 | 0.829±0.004 | 0.853±0.003 |

| | | | | | |
|---|---|---|---|---|---|
| MAERM | **0.984±0.000** | 0.984±0.001 | **0.999±0.000** | **0.999±0.000** | **0.984±0.000** |

**Table S6** Performance comparison of different models on the MAERM-DB from different data sources

| | Acc | Recall | AUC | AUPR | F1 |
|---|---|---|---|---|---|
| **ESP-ER** | | | | | |
| MCAnet | 0.973±0.002 | 0.974±0.003 | 0.996±0.000 | 0.995±0.001 | 0.969±0.002 |
| ML-DTI | 0.903±0.008 | 0.922±0.015 | 0.972±0.003 | 0.973±0.003 | 0.903±0.007 |
| SPEPP | 0.844±0.001 | 0.814±0.001 | 0.933±0.001 | 0.937±0.000 | 0.839±0.001 |
| ESP | 0.889±0.000 | **0.990±0.000** | 0.975±0.000 | 0.976±0.000 | 0.900±0.000 |
| MAARDTI | 0.974±0.001 | 0.974±0.001 | 0.995±0.000 | 0.995±0.000 | 0.975±0.001 |
| HyperAttention DTI | 0.958±0.003 | 0.951±0.002 | 0.993±0.001 | 0.993±0.001 | 0.958±0.003 |
| Rep-ConvDTI | 0.847±0.017 | 0.972±0.004 | 0.902±0.013 | 0.850±0.020 | 0.864±0.012 |
| CACLENS-TU | 0.838±0.003 | 0.952±0.001 | 0.887±0.004 | 0.829±0.003 | 0.855±0.003 |
| MAERM | **0.986±0.000** | 0.985±0.001 | **0.999±0.000** | **0.999±0.000** | **0.986±0.000** |
| **M-CSA** | | | | | |
| MCAnet | 0.651±0.011 | 0.618±0.052 | 0.696±0.024 | 0.751±0.037 | 0.668±0.025 |
| ML-DTI | 0.565±0.033 | 0.569±0.020 | 0.613±0.029 | 0.619±0.027 | 0.570±0.025 |
| SPEPP | 0.635±0.003 | 0.583±0.007 | 0.727±0.001 | 0.708±0.018 | 0.615±0.001 |
| ESP | 0.533±0.018 | 0.771±0.000 | 0.517±0.006 | 0.537±0.015 | 0.625±0.009 |
| MAARDTI | 0.682±0.019 | 0.653±0.026 | 0.758±0.011 | **0.897±0.010** | 0.755±0.018 |
| HyperAttention DTI | 0.611±0.037 | 0.569±0.069 | 0.684±0.022 | 0.671±0.034 | 0.595±0.051 |
| Rep-ConvDTI | 0.752±0.012 | 0.858±0.045 | 0.803±0.014 | 0.746±0.037 | 0.776±0.010 |
| CACLENS-TU | 0.701±0.024 | 0.784±0.057 | 0.740±0.012 | 0.694±0.015 | 0.723±0.030 |
| MAERM | **0.833±0.010** | **0.861±0.020** | **0.896±0.004** | 0.883±0.002 | **0.843±0.010** |

| **RDB** | | | | | |
|---|---|---|---|---|---|
| MCAnet | 0.816±0.008 | 0.818±0.009 | 0.905±0.002 | 0.946±0.004 | 0.855±0.007 |
| ML-DTI | 0.819±0.013 | 0.799±0.033 | 0.913±0.007 | 0.952±0.006 | 0.852±0.014 |
| SPEPP | 0.877±0.017 | 0.842±0.015 | 0.953±0.001 | 0.952±0.003 | 0.872±0.018 |
| ESP | 0.839±0.003 | 0.849±0.007 | 0.922±0.002 | 0.948±0.002 | 0.873±0.003 |
| MAARDTI | 0.758±0.096 | 0.781±0.166 | 0.802±0.064 | 0.878±0.050 | 0.807±0.099 |
| HyperAttention DTI | 0.830±0.003 | 0.824±0.006 | 0.903±0.007 | 0.934±0.009 | 0.864±0.003 |
| Rep-ConvDTI | 0.824±0.032 | 0.960±0.005 | 0.871±0.044 | 0.780±0.082 | 0.846±0.024 |
| CACLENS-TU | 0.780±0.045 | 0.958±0.002 | 0.878±0.048 | 0.858±0.055 | 0.815±0.031 |
| MAERM | **0.930±0.002** | **0.987±0.006** | **0.984±0.000** | **0.992±0.000** | **0.949±0.002** |

**Table S7** Performance comparison of different models on the BioCat-547 dataset

| | Balanced Acc | AUC | AUPR | MCC | Macro-F1 |
|---|---|---|---|---|---|
| MCANet | 0.517±0.020 | 0.550±0.058 | 0.421±0.032 | 0.114±0.050 | 0.393±0.003 |
| MAARDTI | 0.486±0.022 | 0.488±0.020 | 0.410±0.099 | 0.096±0.042 | 0.396±0.010 |
| MAERM | **0.697±0.001** | **0.775±0.002** | **0.655±0.004** | **0.415±0.010** | **0.704±0.003** |

### Part 7. Supplementary Figures

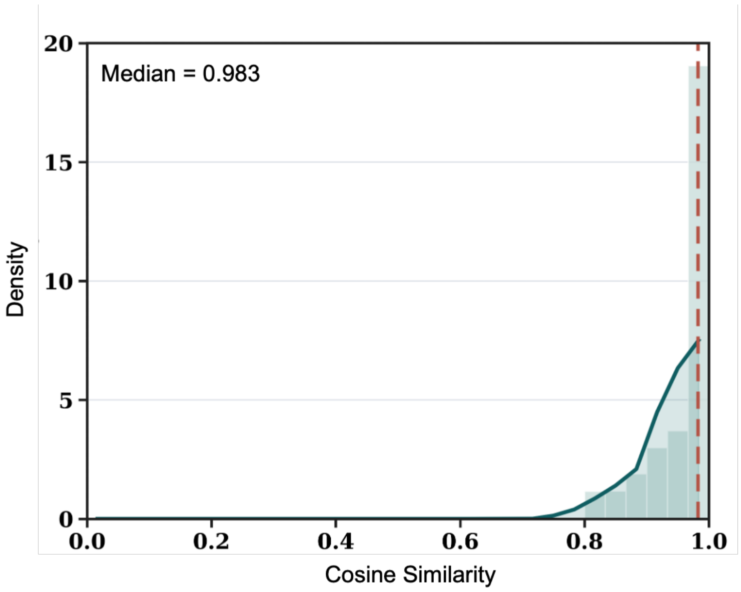


**Figure S1 Histogram and smoothed density** of similarity between the selected negative reactions and positive reactions, with the dashed line marking the median score. Similarity was calculated using rxnfp embeddings [19].

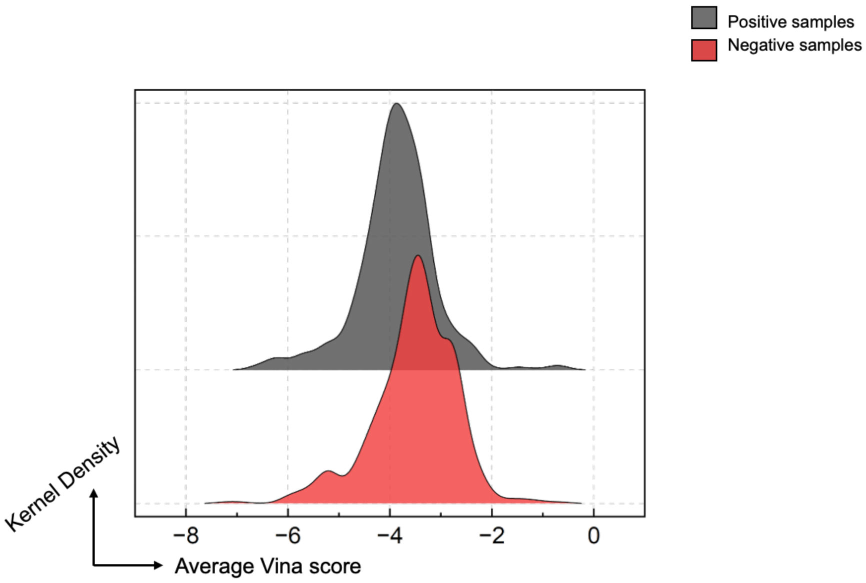


**Figure S2** Ridgeline plot comparing the distributions of average Vina scores for positive and negative enzyme-substrate pairs in the RDB dataset. Only pairs with Vina scores < 0 are shown.

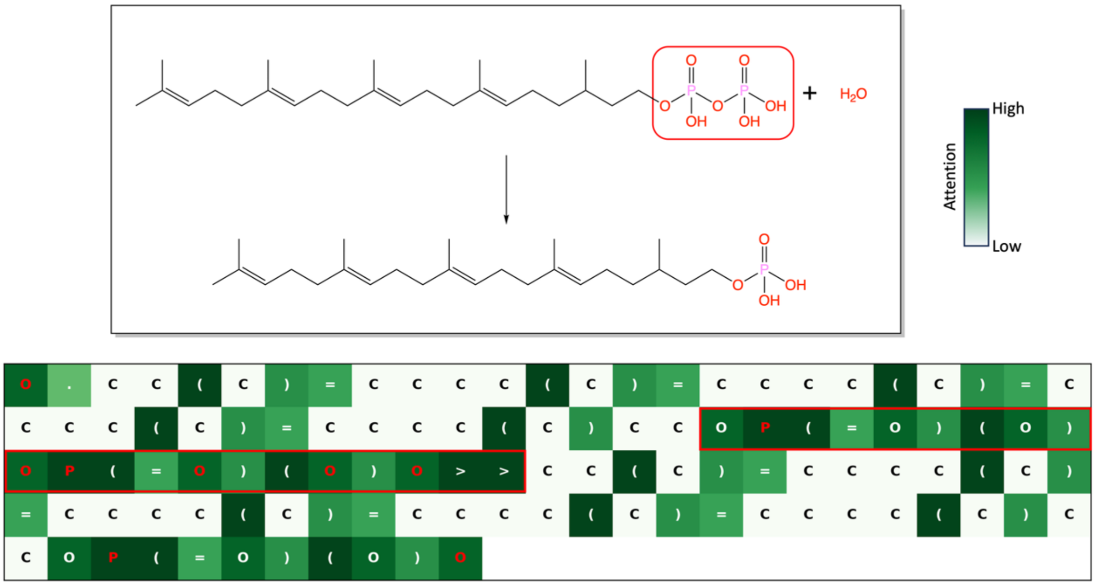


**Figure S3 Attention map for diphosphonic acid hydrolysis.** Red characters indicate the reaction center (radius = 1), and the white-to-green scale denotes increasing attention. The red-boxed region corresponds to the phosphate bond.

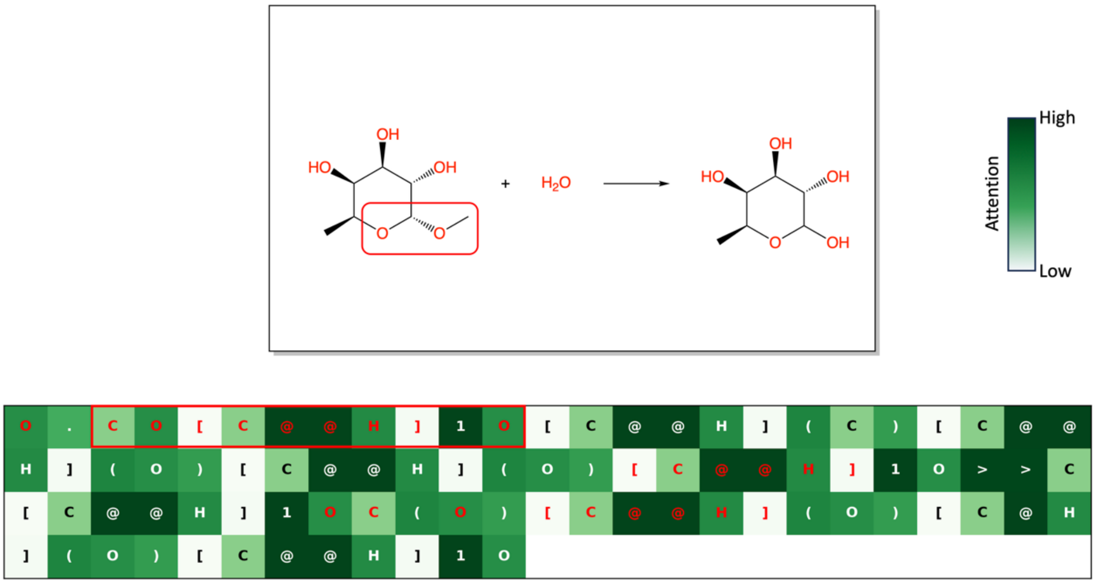


**Figure S4 Attention map for a hydrolysis reaction.** Red characters indicate the reaction center (radius = 1), and the white-to-green scale denotes increasing attention. The red-boxed region corresponds to the glycosidic bond and neighboring atoms.

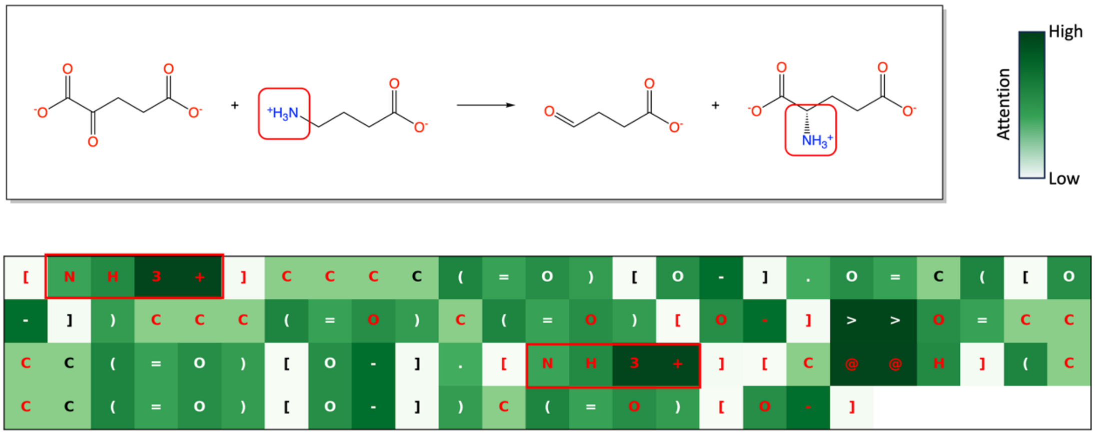


**Figure S5 Attention map for a transamination reaction.** Red characters indicate the reaction center (radius = 1), and the white-to-green scale denotes increasing attention. The red-boxed region corresponds to the amino group.

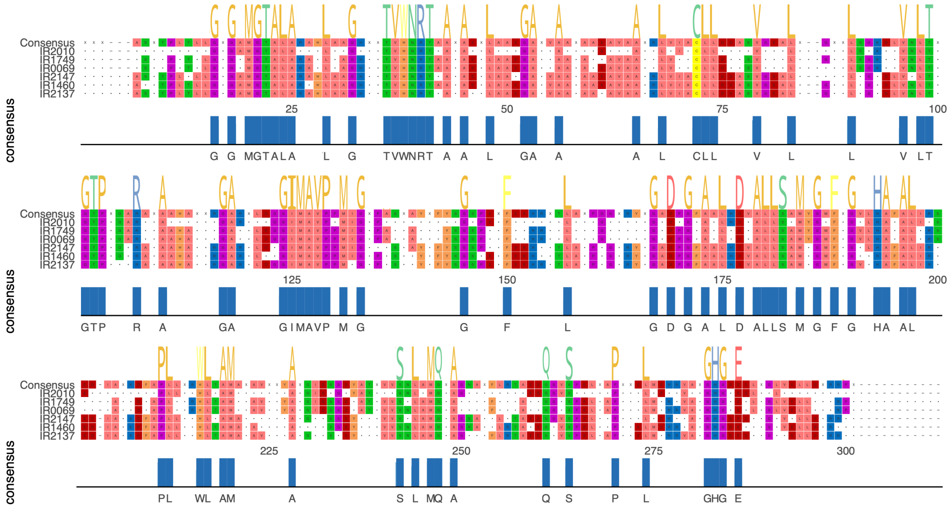


**Figure S6** Sequence similarity analysis of six representative imine reductase (IRED) variants in BioCat-547 with conversion rates above 99%.

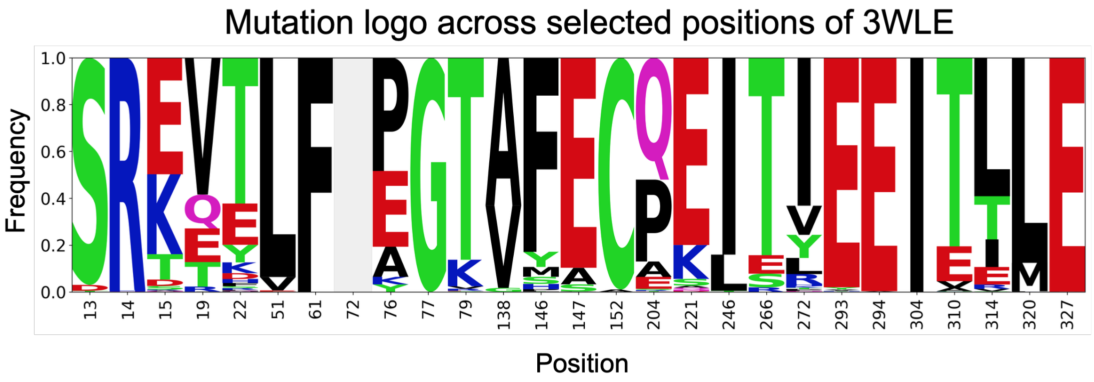


**Figure S7** Mutation logo of selected positions in 3WLE.
